# Resolving the spatial and cellular architecture of lung adenocarcinoma by multi-region single-cell sequencing

**DOI:** 10.1101/2020.09.04.283739

**Authors:** Ansam Sinjab, Guangchun Han, Warapen Treekitkarnmongkol, Kieko Hara, Patrick Brennan, Minghao Dang, Dapeng Hao, Ruiping Wang, Enyu Dai, Hitoshi Dejima, Jiexin Zhang, Elena Bogatenkova, Beatriz Sanchez-Espiridion, Kyle Chang, Danielle R. Little, Samer Bazzi, Linh Tran, Kostyantyn Krysan, Carmen Behrens, Dzifa Duose, Edwin R. Parra, Maria Gabriela Raso, Luisa M. Solis, Junya Fukuoka, Jianjun Zhang, Boris Sepesi, Tina Cascone, Lauren Byers, Don L. Gibbons, Jichao Chen, Seyed Javad Moghaddam, Edwin J. Ostrin, Daniel G. Rosen, John V. Heymach, Paul Scheet, Steven Dubinett, Junya Fujimoto, Ignacio I. Wistuba, Christopher S. Stevenson, Avrum E. Spira, Linghua Wang, Humam Kadara

## Abstract

Little is known of the geospatial architecture of individual cell populations in lung adenocarcinoma (LUAD) evolution. Here, we perform single-cell RNA sequencing of 186,916 cells from 5 early-stage LUADs and 14 multi-region normal lung tissues of defined spatial proximities from the tumors. We show that cellular lineages, states, and transcriptomic features geospatially evolve across normal regions to LUADs. LUADs also exhibit pronounced intratumor cell heterogeneity within single sites and transcriptional lineage-plasticity programs. T regulatory cell phenotypes are increased in normal tissues with proximity to LUAD, in contrast to diminished signatures and fractions of cytotoxic CD8+ T cells, antigen-presenting macrophages and inflammatory dendritic cells. We further find that the LUAD ligand-receptor interactome harbors increased expression of epithelial CD24 which mediates pro-tumor phenotypes. These data provide a spatial atlas of LUAD evolution, and a resource for identification of targets for its treatment.

**Statement of significance:** The geospatial ecosystem of the peripheral lung and early-stage LUAD is not known. Our multi-region single-cell sequencing analyses unravel cell populations, states, and phenotypes in the spatial and ecological evolution of LUAD from the lung that comprise high-potential targets for early interception.

## INTRODUCTION

Lung adenocarcinoma (LUAD) is the most common histological subtype of lung cancer and accounts for most cancer deaths (1,2). Over the past decade, annual low dose CT screening was endorsed in an effort to reduce lung cancer mortality (3). Since then, an increasing number of early-stage LUAD diagnoses has warranted the need for novel personalized early treatment strategies. This in turn heavily rests on improved understanding of molecular and cellular processes underlying early LUAD development.

Previous studies identified molecular alterations in histologically normal-appearing epithelial *fields* that are close to solid tumors including those of the lung and that are less prevalent or absent in relatively more distant (from the tumor) regions -- suggesting geospatial heterogeneity in the uninvolved lung that is pertinent to development of a nearby tumor (4). While these studies provided valuable insights into the *spatial* development of cancer from a particular niche in the lung, they have been mainly guided by bulk profiling approaches (4,5). It is now appreciated that editing of the immune microenvironment towards protumor phenotypes including escape of immune surveillance portends the underlying biology, development, and progression of LUAD (5). Yet, the interplay between individual immune cell populations and other cell subsets in geospatial development of early-stage LUAD is not known. Technologies that profile tissues at single-cell resolution have permitted delineating the molecular and cellular complexity of tumor ecosystems. Single-cell sequencing technologies were used to chart the immune microenvironment in metastasis and therapeutic response of advanced lung cancers to targeted therapies (6–9). Yet, the complex spatial evolution of heterogeneous cellular populations and their interactions, and as an early-stage LUAD develops from the peripheral lung, remains largely unresolved.

Here, we sought to discern the spatial atlas of the peripheral lung and early-stage LUAD at single-cell resolution to better understand the topological architecture of LUAD evolution. We performed deep scRNA-seq analysis of 19 spatial regions, including enriched epithelial populations, from 5 early-stage LUADs and 14 multi-region normal-appearing lung tissues with differential and defined spatial proximities from the tumors. Our study unravels tumor evolutionary trajectories as well as geospatial evolution in cell populations and their expression signatures that portray how early-stage LUAD develops from the lung ecosystem.

## RESULTS

### Single-cell spatial landscape of early-stage LUAD

To begin to chart a comprehensive single-cell atlas of early-stage LUAD and the peripheral lung, we performed scRNA-seq on all cells from an early-stage LUAD (P1, **Supplementary Table S1**) as well as matched tumor-adjacent and relatively more distant normal lung tissues (**Fig. 1A**). Unsupervised clustering of 15,132 QC-passed cells revealed cell clusters representing 5 major cellular lineages, namely epithelial, endothelial, myeloid, lymphoid, and stromal cell subsets (**Fig. 1B**, **Supplementary Fig. S1**). Epithelial (*EPCAM*+) cell fractions were 3.7%, 5.4%, and 3.5% for tumor, tumor-adjacent and -distant normal samples, respectively, at an average of 4.2% and in line with previous studies ((8); **Fig. 1B-C**).

**Figure 1.**
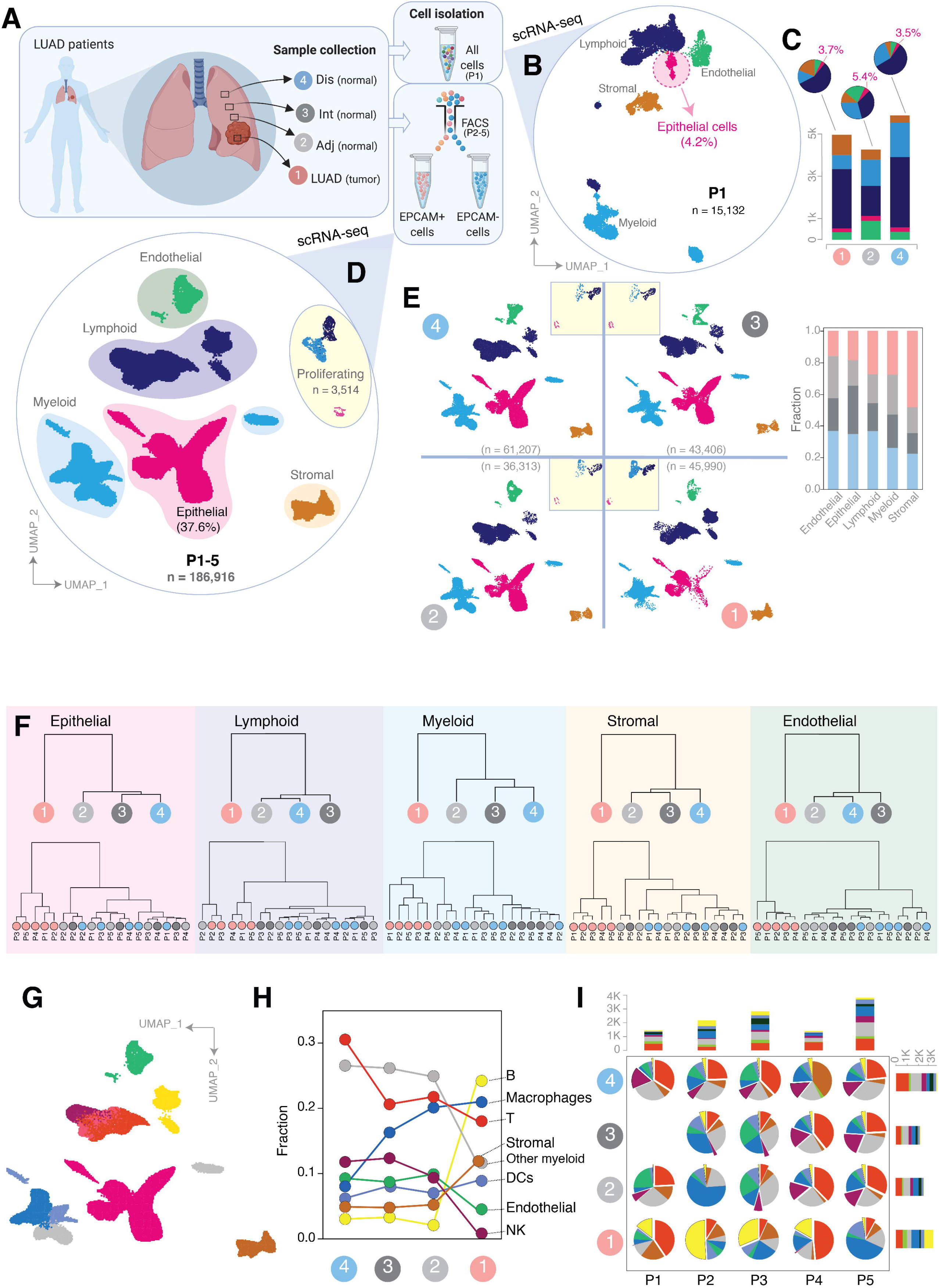
Dissecting early-stage LUAD and the peripheral lung ecosystem by multi-region single-cell RNA sequencing. **A**, Workflow showing multi-region sampling strategy of 5 LUADs and 14 spatially defined normal lung tissues for analysis by scRNA-seq. Dis, distant normal; Int, intermediate normal; Adj, adjacent normal; LUAD, tumor tissue. **B**, Uniform manifold approximation and projection (UMAP) embedding of cells from tumor, adjacent normal and distant normal samples of patient one (P1). Cells are colored by their inferred cell types. **C**, Cell composition in absolute cell numbers (stacked bar plots) and relative fractions (pie charts) in each spatial sample derived from P1. **D**, UMAP view of cells from all 5 patients, including EPCAM+ and EPCAM-pre-enriched cells from P2-P5. Colors represent assigned major cell types. Proliferating, proliferating cells. **E**, UMAP view of cell types and their fractions (stacked bar plot) by spatial samples. Clusters in embedded squares represent proliferating cells. Colors represent assigned cell types as in panel D. **F**, Dendrograms showing hierarchical relationships of cells among the spatial samples based on the computed Euclidean distance using transcriptomic features. Dendrograms are shown for 5 major cell types (from left to right), for all patients together (top) and by patient (bottom). **G**, Same UMAP as in panel D, with further subclustering of lymphoid and myeloid cells. Colors correspond to the cell type annotation in panel H for EPCAM-cells. **H-I**, Line plot showing changes in the relative fractions among the EPCAM negative subsets across spatial samples for all patients together (H) and by patient (pie charts, I). Stacked bar plots in panel I show absolute cell numbers of the fractions by patient and spatial sample.

To increase the throughput and to better capture patterns of cellular heterogeneity based on distance from LUADs, in particular within the epithelial lineage, we performed separate scRNA-seq analysis of epithelial (*EPCAM*+) and non-epithelial (*EPCAM*-) cells enriched from early-stage LUADs of four additional early-stage LUAD patients (P2-P5, Methods, **Supplementary Fig. S1; Supplementary Table S1**), each with three matching normal lung tissues of defined spatial proximities to LUADs: tumor-adjacent, - intermediate and -distant (19 samples and 35 scRNA-seq libraries from P1-P5; **Fig. 1A**). The spatial locations of multi-region normal tissues were carefully defined with respect to the tumor edge (see Methods), such that the samples span a spatial continuum and, thus, enable interrogation of geospatial relationships among early-stage LUAD and the peripheral lung tissues. A total of 186,916 cells, uniformly derived from all patients and sequencing batches, were retained for subsequent analyses with a median of 1,844 detected genes per cell (**Supplementary Fig. S2A-C; Supplementary Table S1**). Cells clustered into the above-mentioned 5 distinct lineages (**Supplementary Fig. S2D**), and clustering was deemed robust based on high ratio of overlapping cell memberships with an independent clustering method (reciprocal PCA), and with clusters of down-sampled cells (**Supplementary Fig. S3A-B**; Supplementary Methods).

We were able to profile samples markedly enriched with epithelial cells (37.6%, n = 70,030 epithelial cells, including 3,514 proliferating cells) in comparison to the unbiased approach in P1 (4.2%) (**Fig. 1D**). Cells were adequately derived from all spatial samples and their lineage compositions varied spatially across the LUAD and normal tissues (**Fig. 1E**). Proliferating epithelial cells were highly enriched in LUADs (40%) compared to normal tissues (*P* = 0.07; **Supplementary Fig. S2E**). We also noted higher transcriptome complexity (number of genes) in EPCAM+ fractions compared to EPCAM-fractions from P2-P5 (**Supplementary Fig. S2A** right; **Supplementary Table S1**). This was also significantly evident among cells of P1 which were not subjected to enrichment by EPCAM-based cell sorting (**Supplementary Fig. S4A-F**; *P* < 0.01).

Next, we analyzed hierarchical relationships among major cell lineages and found that cells from all patient LUADs were transcriptomically distinct from their matched spatial normal counterparts (**Fig. 1F**, see Methods). Overall, cells from adjacent normal samples clustered more closely with those of the LUADs than with more distant normal tissues (e.g., epithelial and myeloid lineages, **Fig. 1F** top dendrograms). This average pattern of transcriptomic similarity between tumor and adjacent normal samples was evident in specific patients (e.g., P2, **Fig. 1F** bottom dendrograms). We then further classified lymphoid and myeloid lineages into major cell types (**Fig. 1G**; **Supplementary Fig. S2F,** Methods). Analysis of spatial cell composition revealed distinct topological gradients with greater tumor proximity, which were consistently evident across patients, as well as increased fractions of B cells and decreased abundance levels of NK cells specifically in the LUADs (**Fig. 1H-I**; **Supplementary Table S2**). These observations highlight geospatial transcriptomic heterogeneity in tumor microenvironment (TME) landscape of early-stage LUADs.

### Spatial diversity of lung epithelial cells and intratumoral heterogeneity in LUAD

We next interrogated spatial epithelial features of the LUADs and multi-region normal tissues. The 70,030 epithelial cells derived from all samples clustered into 10 distinct epithelial cell types with high degree of robustness (**Fig. 2A**, **Supplementary Fig. S5A-C**). These clusters represented alveolar type I (AT1; C2, *AGER*+), AT2 (C3; *SFTPC*+), basal (C4; *KRT15*+), bronchioalveolar (C5; *SFTPC*+/*SCGB1A1*+), ciliated (C6; *PIFO*+) and club/secretory (C7; *BPIFB1*+) cells (**Fig. 2A-B**; **Supplementary Table S3**). We also identified the recently described and rare ionocytes (C8; *FOXI1*+/*CFTR+*; (10,11)), bipotent alveolar progenitors (C1; (12)), and unique cell states such as proliferating basal cells (C10; *TOP2A+*) (**Fig. 2A-B**; **Supplementary Table S3**).

**Figure 2.**
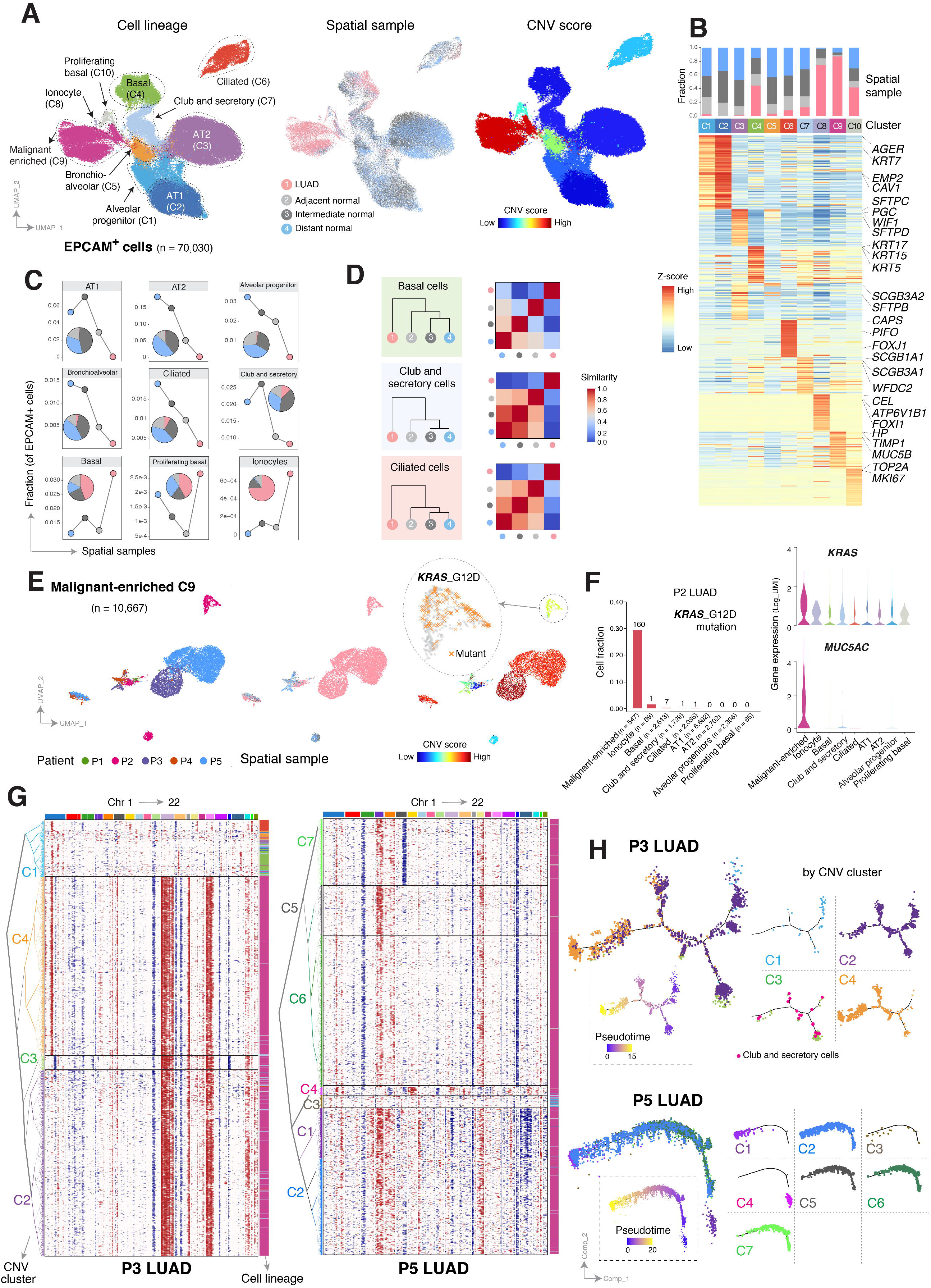
Epithelial lineage diversity and intratumoral heterogeneity in the spatial ecosystem of early-stage LUAD. **A**, UMAP visualization of all EPCAM+ cells from P1-P5 colored (from left to right) by their assigned cell types, spatial samples, and inferred copy number variation (CNV) scores. **B**, Heatmap of major lineage marker genes for EPCAM+ cell clusters (C1-10 as shown in panel A, left), with corresponding bar blots outlining fraction by spatial sample. **C**, Area plot showing changes in EPCAM+ subset fractions across spatial samples. **D**, Hierarchical relationships of 3 representative subsets of epithelial cells (from top to bottom) among the spatial samples based on the computed Euclidean distance using transcriptomic features (left), and corresponding heatmaps quantifying similarity levels among spatial samples (right). Similarity score is defined as one minus the Euclidean distance. **E**, UMAP plots of cells in the malignant-enriched cluster C9 (panel A), colored by their corresponding patient origin (left), spatial sample (middle), and CNV score (right). The zoom in view of the right panel shows *KRAS* G12D mutant cells in P2. **F**, Fraction of cells carrying *KRAS* G12D mutation (left bar plot), with numbers indicating the absolute cell numbers, as well as expression levels of *KRAS* (violin plot, top right) and *MUC5AC* (violin plot, bottom right), within cells of each epithelial lineage cluster of P2. **G**, Unsupervised clustering of CNV profiles inferred from scRNA-seq data from patient P3 (left) and P5 (right) tumor samples using NK cells as control and demonstrating intratumoral heterogeneity in CNV profiles. Chromosomal amplifications (red) and deletions (blue) are inferred for all 22 chromosomes (color bars on the top). Each row represents a single cell, with corresponding cell type annotated on the right (same as in panel A). **H**, Potential developmental trajectories for EPCAM+ cells from P3 (top) and P5 (bottom) inferred by Monocle 3 analysis. Cells on the tree are colored by pseudotime (dotted boxes) and CNV clusters.

We noted a malignant-enriched cluster (C9) with cells of mixed lineages (8) from all patients, mostly their LUADs (**Fig. 2A-B**, **Supplementary Fig. S6A-B**). Interestingly, few cells from the normal tissues were also found in the C9 cluster (**Fig. 2B**, **Supplementary Fig. S6A**). To distinguish *bona fide* malignant cells from non-malignant subsets, we employed a strategy that infers copy number variations (CNVs) from scRNA-seq data in every epithelial cell and we generated a CNV score to quantify their level of aneuploidy (13) (see Supplementary Methods). Cells in C9 exhibited overall increased CNV scores as well as higher amplitudes of CNVs (**Fig. 2A** right), thereby supporting the overall malignant assignment of this cluster. While cells from the LUADs predominantly resided in C9, a fraction (29%) was transcriptomically and genotypically (e.g., reduced copy number profiles) similar to basal cells (17%; **Supplementary Fig. S6C; Supplementary Table S4**). Interestingly, basal cells were enriched in all LUADs compared to their normal samples (*P* = 0.09; **Supplementary Fig. S6B and D; Supplementary Table S5**). We also noted pronounced steady-state enrichment or depletion of epithelial subsets with spatial proximity to the tumors (**Fig. 2C**). Relative to cells from tumor-intermediate or -distant normal sites, cells from tumor-adjacent normal tissues were, overall, more transcriptomically similar to (clustered closely with) those from the LUADs (**Fig. 2D**).

Alveolar differentiation hierarchies have been shown to partake in lung tumor development *in vivo* (12,14,15). In our cohort, alveolar cells with definitive lineage features (e.g., AT1, AT2, and alveolar progenitors) were depleted in LUAD tissues (**Fig. 2C**), which prompted us to leverage the relatively large number of alveolar cells sequenced to dissect potential alveolar differentiation trajectories. Pseudotemporal ordering of alveolar cells revealed a developmental hierarchy that was initiated by AT2 cells and that followed a main trajectory of differentiation into AT1 cells (**Supplementary Fig. S7A-B**) in close agreement with previous studies in mice (12,14,15). Given the reported role of NOTCH in AT2-to-AT1 differentiation and alveolar repair (16), we interrogated a NOTCH signaling score which we found to be increased along the alveolar differentiation trajectory (**Supplementary Fig. S7C-D**).

To further investigate malignant programs, we performed subclustering of cells from malignant-enriched cluster C9 (n = 10,667) while overlaying CNV scores, which separated likely malignant cells from subsets derived from normal tissues (**Fig. 2E**, **Supplementary Fig. S8A-E; Supplementary Table S6**). We noted low CNV scores (**Supplementary Fig. 8A** and **Fig. 2E** right) in cells of the malignant-enriched cluster in the LUAD of P2. We thus interrogated the presence of *KRAS* codon 12 driver mutations which reside within genomic brackets captured in our 5’ single-cell sequencing design (see Supplementary Methods). Interestingly, among all P2 epithelial subsets, 29% of C9 cells (160 of 547) harbored the *KRAS* G12D mutation (**Fig. 2E** right **and 2F**) and were exclusively derived from the tumor sample. Compared to *KRAS* wild-type cells from the same LUAD, cells harboring *KRAS* G12D mutation exhibited distinctively high expression of tumor markers (e.g., *CEACAM5*; **Supplementary Fig. S8B**), increased expression of *MUC5AC* (**Fig. 2F**) and *LCN2* as well as reduced expression of *NKX2-1* (**Supplementary Fig. S8B; Supplementary Table S7**), altogether suggestive of mucinous differentiation (17,18) and in line with the histological (mucinous) pattern of this tumor (**Supplementary Table S1**). These findings underscore spatial heterogeneity patterns in *KRAS* driver mutation and cellular lineage that are likely unique to the ecosystem of *KRAS*-mutant LUAD. Additional clustering of tumor-derived C9 cells by patient underscored transcriptomic features that were shared between 2 or more LUADs (e.g., increased *CEACAM5* or *CEACAM6*) (**Supplementary Fig. S8B-E**). We also noted patient/LUAD-specific transcriptomic features such as enrichment of club and secretory (P2) or AT2 (P5) canonical markers thus potentially signifying distinct cells-of-origin among the LUADs (**Supplementary Fig. S8B and E**).

Unlike P2, C9 cells in P3 and P5 were almost exclusively derived from the LUAD tissues (**Fig. 2E**). In P3 LUAD, we identified large-scale chromosomal alterations (**Fig. 2G** left, **Supplementary Fig. S9A; Supplementary Table S6**), based on which unsupervised clustering analysis revealed 4 clusters with differential CNV profiles. Among them, three clusters (C2, C3, and C4) exhibited pronounced CNVs that were indicative of malignant cell features (**Fig. 2G** left). We found an additional CNV event (i.e., gain of 1p) unique to cells of cluster C4 but not C2 or C3, possibly signifying a late event in the evolutionary trajectory of P3 LUAD. We also observed a branched trajectory that started with cells of C2 and C3 and comprised few “normal cells” with club/secretory lineage, and that later branched into cells of CNV cluster C4 (**Fig. 2H** top) -- suggesting that C4 evolved from C2 and C3 and that P3 LUAD perhaps originated from club/secretory cells. This is consistent with increased expression of club/secretory canonical markers (e.g., *TFF3, HP, MUC4*) in P3 tumor clones with also high CNV scores (C0 and C2; **Supplementary Fig. S8C**). P5 LUAD comprised 7 distinct CNV clusters (**Fig. 2G** right, **Supplementary Fig. S9B**). C1 harbored cells with increased CNV events, including events with higher amplitude, while C4 clustered the closest to C3 which mostly comprised non-malignant cells (**Fig. 2G** right). Pseudotime analysis revealed a C4-to-C1 path which, in contrast to P3, was unbranched suggesting that C4 and C1 in P5 likely comprised malignant cells from early and late developmental states, respectively (**Fig. 2H** bottom). Overall, hierarchical clustering analysis was consistent with phylogenetic reconstruction of tumor clonal architecture using inferred large-scale CNVs (*r* = 0.7 in P3; **Supplementary Fig. S10**). Our single-cell interrogation of a large number of epithelial cells from multi-region tissues identified diverse epithelial identities, malignant trajectories, as well as high-resolution intratumor cell heterogeneity.

### Lymphoid reprogramming towards a protumor microenvironment

We further characterized lymphoid spatial dynamics (**Fig. 1H-I**) and states (n = 53,882 cells, see Methods). Following clustering, we found 10 transcriptomically distinct lymphoid cell types/states that were uniformly derived from all sequencing batches and patients (**Fig. 3A**, **Supplementary Fig. S11A-B; Supplementary Table S8**). Lymphoid clusters were overall spatially modulated by tumor proximity (**Fig. 3C**). Relative to normal tissues, LUADs were heavily enriched with plasma *(SDC1+/MZB1+)*, B (*CD19+/CD22+*), and T regulatory (Treg; *FOXP3+*) cells (**Fig. 3C**, **Supplementary Fig. S11C**). With increasing tumor proximity, we noted a gradual decrease in NK cells (*GNLY+*), innate lymphoid cells (ILCs), both *GZMA*-hi and *GNLY*-hi CD4+ cytotoxic T lymphocytes (CTL; *CD40LG+, BATF+*), and *GNLY-*hi CD8+ CTLs, all of which were, overall, depleted in the LUADs (**Fig. 3C**, **Supplementary Fig. S11C**).

**Figure 3.**
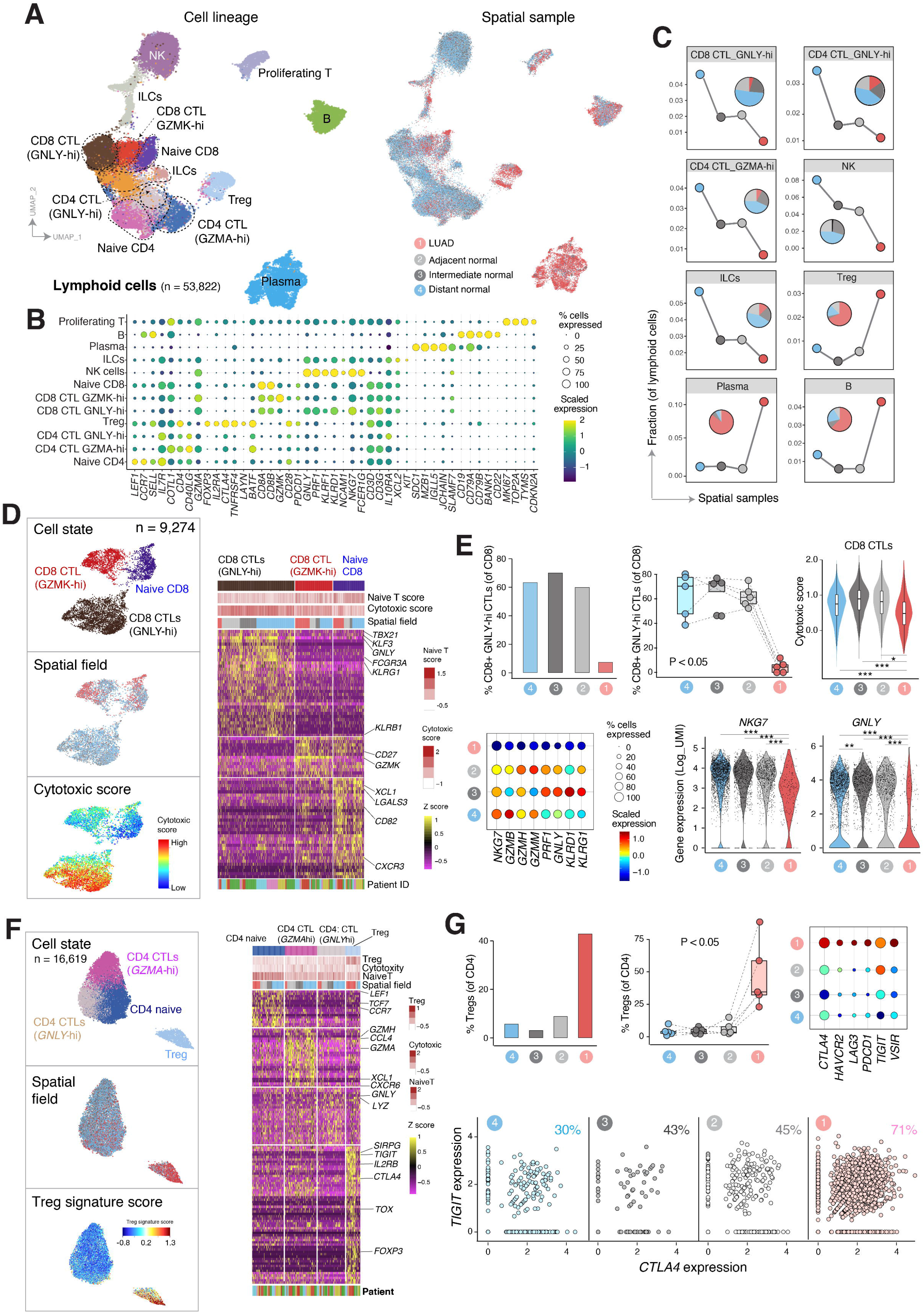
Spatial reprogramming of lymphoid subsets towards protumor phenotypes in early-stage LUAD. **A**, UMAP visualization of lymphoid cell subsets from P1-P5 colored by cell lineage (left) and spatial sample (right). CTL, cytotoxic T lymphocyte; Treg, T regulatory cell; ILC, innate lymphoid cell; NK, natural killer cell. **B**, Bubble plot showing the expression of lineage markers. Both the fraction of cells expressing signature genes (indicated by the size of the circle) as well as their scaled expression levels (indicated by the color of the circle) are shown. **C**, Changes in the abundance of lymphoid cell lineages and cellular states across the LUADs and spatial normal samples. Embedded pie charts show the contribution of each spatial sample to the indicated cell subtype/state. **D**, UMAP plots of CD8+ T lymphocytes colored by cell states (top), spatial sample (middle), and cytotoxic score (bottom). The heatmap on the right shows normalized expression of marker genes for defined CD8 T cell subsets. Each column represents a cell. Top annotation tracks indicate (from top to bottom) cell states, naïve T cell scores and cytotoxic scores calculated using curated gene signatures, and the corresponding spatial sample of each cell. **E**, Depletion of CD8+ GNLY-hi CTLs in the tumor microenvironment (TME) of LUADs. Bar plot (top left) and boxplot (top middle) showing percentage of CD8+ GNLY-hi CTLs among total CD8+ cells from all patients across the spatial sample. Each circle in the boxplot represents a patient sample. *P –* value was calculated using Kruskal-Wallis test. Cytotoxicity signature score (violin plot, top right) of CD8+ CTLs across spatial samples (*, *P* < 0.05; ***, *P* < 0.001). *P –* values were calculated using Wilcoxon rank sum test. The percentage of CD8+ CTLs expressing cytotoxic signature genes (indicated by the size of the circle) and their scaled expression levels (indicated by the color of the circle) across the LUADs and spatial normal lung samples (bubble plot, bottom left). Expression levels of *NKG7* and *GNLY* in CD8+ CTLs across the spatial samples (violin plots, bottom right; **, *P* < 0.01; ***, *P* < 0.001). *P* – values were calculated using Wilcoxon rank sum test. **F**, UMAP plots of CD4+ T lymphocytes colored by cell states (top), spatial sample (middle), and Treg signature score (bottom). The heatmap on the right shows normalized expression of marker genes for CD4+ T cells grouped by defined subcluster. Each column represents a cell. Top annotation tracks indicate (from top to bottom) cell states, Treg signature score, cytotoxic scores, and naïve T cell score calculated using curated gene signatures, and the corresponding spatial sample of each cell. **G**, Enrichment of CD4+ T regulatory cells (Treg) in the TME of LUADs. Bar plot (top left) and boxplot (top middle) showing percentage of CD4+ Tregs among total CD4+ cells from all patients across the spatial samples. Each circle in the boxplot represents a patient sample. *P* – value was calculated using Kruskal-Wallis test. Percentage of CD4+ Tregs expressing inhibitory immune checkpoint genes (indicated by the size of the circle) and their scaled expression levels (indicated by the color of the circle, color assignment same as panel E) across the spatial samples (bubble plot, top right). Frequency of CD4+ Treg cells co-expressing *CTLA4* and *TIGIT* immune checkpoints across the spatial samples (scatter plots, bottom). The fractions of *CTLA4*+*TIGIT*+ Tregs are labeled on each plot.

We further performed subclustering analysis of CD8+ T cells which identified 3 robust clusters: naïve, *GZMK*-hi, and *GNLY*-hi subpopulations with differentially expressed cell state signatures (**Fig. 3D**, **Supplementary Fig. S12; Supplementary Tables S9 and S10**). Consistently, naïve CD8+ T cells showed high naïve and low cytotoxic T cell scores and were composed of cells across all samples in the LUAD space (**Fig. 3D**, **Supplementary Fig. S13A**). In contrast, *GNLY*-hi CD8+ CTLs exhibited high levels of cytotoxicity genes (*TBX21, KLF3, FCGR3A, KLRG1*, *KLRB1*) and scores and were depleted in the LUAD samples (**Fig. 3C-E**). We observed a significant spatial pattern of reduced cytotoxic activity in P3 and P4 (**Supplementary Fig. S13B** left and right, respectively; *P* < 0.05). Overall, CD8+ CTLs showed significant and spatially-modulated reduction in cytotoxicity signature score (*P* < 0.05) and decreased expression of major cytotoxic genes such as *NKG7* and *GNLY* (*P* < 0.01; **Fig. 3E**).

Spatial analysis of CD4+ T cell states (**Supplementary Tables S10 and S11**) showed that LUAD tissues were specifically enriched with *FOXP3+* Tregs (**Fig. 3F-G; Supplementary Table S12**) and the Treg signature scores were significantly and spatially increased with proximity to all LUADs (**Supplementary Fig. S13C** left; *P* < 0.01) and in each of P3 and P5 (**Supplementary Fig. S13C** middle and right, respectively; *P* < 0.05). The Tregs cells also expressed high levels of pro-tumor immune checkpoints including *TIGIT, CTLA4, LAG3,* or *PDCD1* (**Fig. 3G**). The fraction of Tregs co-expressing both *CTLA-4* and *TIGIT* immune checkpoints was progressively higher along the spectrum of distant normal sites to more adjacent-to-tumor regions up to the LUADs (**Fig. 3G** bottom). In contrast, we noted a reduction of cytotoxic CD4+ CTLs characterized by high expression of *GZMA* (*P* < 0.05), or co-expression of *GZMA* and *GZMH*, with increasing proximity to all LUADs (**Supplementary Fig. S13D-F**).

We further examined the spatial enrichment of LUADs with plasma and B cells (**Fig. 3C; Supplementary Table S12**). We found spatial changes in plasma cell isotype-switching, such as increased *IGHA1/2* and decreased *IGHG1/3*, with increasing proximity to P3 and P5 LUADs (**Supplementary Fig. S13 G-I; Supplementary Table S13**). Smoker patients (P2 and P3) harbored strikingly high plasma cell fractions relative to nonsmokers (P1 and P5) (**Supplementary Fig. S14A**). We confirmed the increased plasma fractions in smoker LUADs following analysis of TCGA cohort (*P* < 0.001; **Supplementary Fig**. **S14B**). We also identified 3 distinct subsets of B cell states (**Supplementary Table S14**), including a LUAD-enriched subcluster (C0) with high expression levels of *RAC2+* and *ACTG*+ (**Supplementary Fig. S13J-L**), known to play key roles in synapse formation in B cells (19). The B cell signature (C0) was progressively increased across atypical adenomatous hyperplasias (AAH), the preneoplastic precursors of LUAD, and invasive LUADs compared to matched normal lung tissues (**Supplementary Fig. S13M**). The B cell signature was also associated with prolonged overall survival (OS) and progression free interval (PFI) in treatment-naïve LUAD patients from TCGA (*P* = 0.0001, OS; *P* = 0.02, PFI; **Supplementary Fig. S15A**) and in-house (MDACC; *P* = 0.2, OS; *P* = 0.03, PFI; **Supplementary Fig. S15B**) cohorts. Our analyses identify spatial properties in lymphoid cell states that may underlie pro-tumor immune remodeling in early-stage LUAD.

### Depletion of antigen presenting macrophages and inflammatory dendritic cells in early-stage LUAD

Spatial myeloid patterns (**Fig. 1H-I**) in LUAD space prompted us to further investigate myeloid subsets and cellular states (**Fig. 4**). In total, 45,803 myeloid cells from all sequencing batches and patients analyzed in this study (**Supplementary Fig. S16A-B**) clustered into 13 distinct subsets: classical monocytes (*S100A8+, S100A9+*), non-classical monocytes *(CDKN1C+*), mast cells *(MS4A2+)*, neutrophils (*IL1A+*), M2-like macrophages C1 (*TREM2+)*, M2-like macrophages C5 (*CD163+*), alveolar macrophages (*MARCO+)*, classical DC 1 (cDC1; *CLEC9A+*), cDC2 (*CLEC10A+*), plasmacytoid DC (pDC; *PLD4+*), other DCs (*CCL22+)* and proliferating myeloid cells *(TOP2A+)* (**Fig. 4A-B**; **Supplementary Tables S15 and S16**). Myeloid clusters were derived from tumor and all normal spatial samples with varying proportions (**Supplementary Fig. S16C**). M2-like macrophages C5, monocytes (classical and non-classical), and mast cells were gradually depleted with increasing tumor proximity, whereas M2-like macrophages C1, proliferating myeloid subsets and cDC2 cells were steadily enriched in the tumors (**Fig. 4A** **and** **C**, **Supplementary Fig. S17A**).

**Figure 4.**
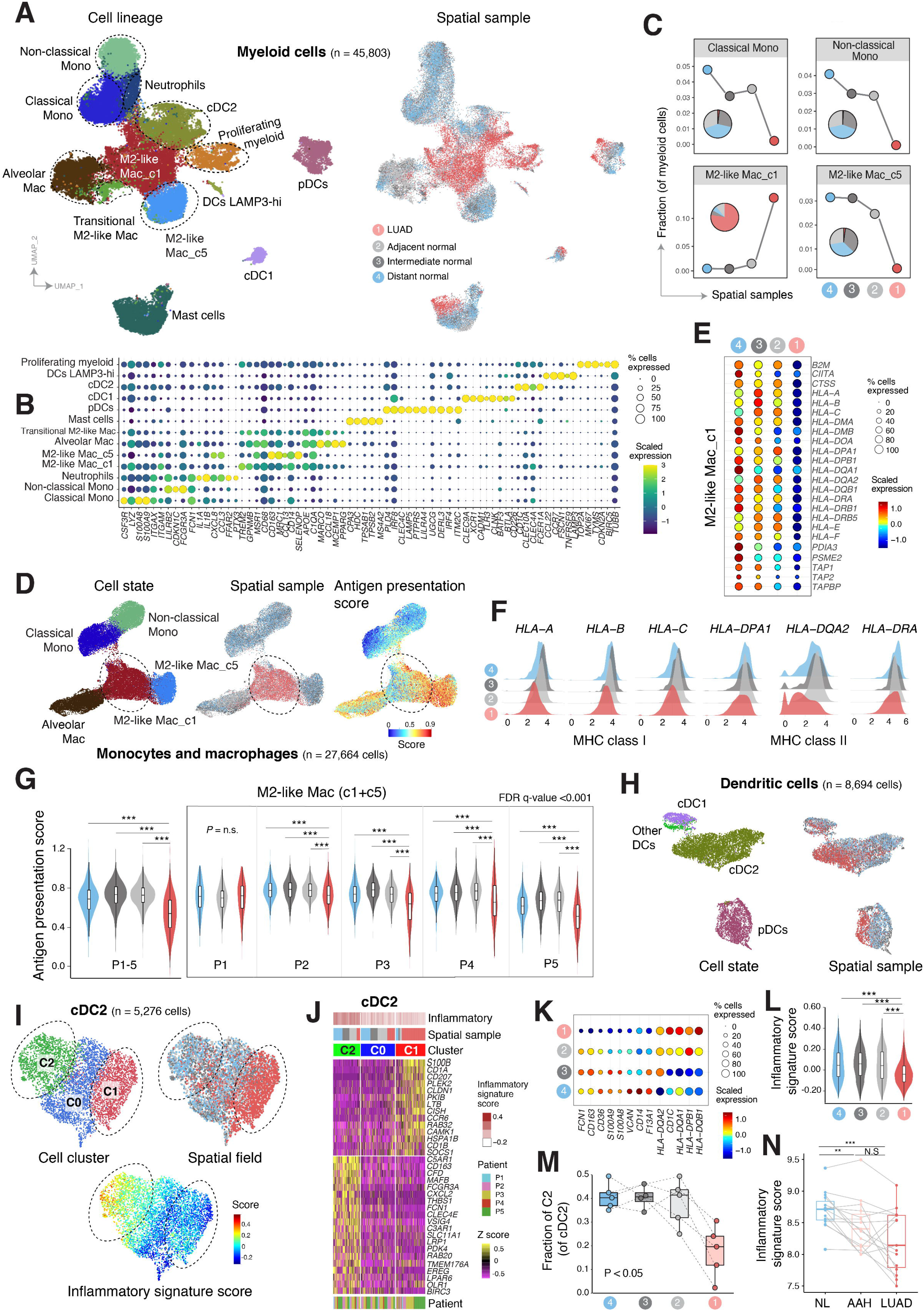
Reduced signatures of antigen presentation and inflammatory dendritic cells in the microenvironment of early-stage LUAD. **A**, UMAP visualization of myeloid cell lineages colored by cell type/state (left) and the spatial samples (right). Mac, macrophages; Mono, monocytes; DC, dendritic cell; cDC, classical dendritic cell; pDC, plasmacytoid dendritic cell. **B**, Bubble plot showing the percentage of myeloid cells expressing lineage specific marker genes (indicated by the size of the circle) as well as their scaled expression levels (indicated by the color of the circle). **C**, Changes in the abundance of myeloid cell subsets across the LUADs and spatial normal lung samples. Embedded pie charts show the contribution of each spatial sample to the indicated cell subtype/state. **D**, UMAP plot of monocyte and macrophage subpopulations, color coded by cell type/state (left), spatial sample (middle), and antigen presentation score (right). **E**, Percentage of M2-like macrophages cluster 1 expressing antigen presentation genes (indicated by the size of the circle) and their scaled expression levels (indicated by the color of the circle) across the spatial samples (bubble plot). **F**, Ridge plots showing the distribution of MHC class I and MHC class II gene expression densities in M2-like macrophages cluster 1, and across LUADs and spatial normal lung samples. **G**, Violin plots showing the antigen presentation score in M2-like macrophages (clusters 1 and 5) across LUADs and spatial normal lung samples for all patients together (left) and within patients (right) (***, *P* < 0.001, N.S, *P* > 0.05). *P* – values were calculated by Wilcoxon rank sum test. **H**, UMAP plots of dendritic cells, color coded by cell state (left) and spatial sample (right). **I**, UMAP plots showing unsupervised subclustering of cDC2 cells colored by cluster ID (top left), spatial sample (top right) and the computed inflammatory signature score (bottom). **J**, Heatmap showing normalized expression of marker genes of cDC2 cell subsets. The top annotation tracks indicate (from top to bottom) the inflammatory signature scores, spatial tissue of origin, and cDC2 cell clusters. **K**, Bubble plot showing the percentage of cDC2 cells expressing inflammatory and non-inflammatory signature genes (indicated by the size of the circle) as well as their scaled expression levels (indicated by the color of the circle) in the LUADs and spatial normal lung tissues. **L**, Violin plot showing expression of inflammatory signature score in cells from cDC2 cluster C2 in the TME of LUADs. *P* – value was calculated using Kruskal-Wallis test (***, *P* < 0.001). **M**, Boxplot showing fraction of cDC2 C2 cells among total cDC2 cells and across the LUADs and spatial normal lung tissues. Individual circles correspond to patient samples. *P* – value was calculated using Kruskal-Wallis test. **N**, Violin plot showing the inflammatory signature scores in cDC2 C2 cells across LUADs and spatial normal lung samples. **O,** boxplot showing the inflammatory signature score in normal lung (NL), in premalignant atypical adenomatous hyperplasia (AAH) and in LUAD from an independent cohort (**, *P* < 0.01; ***, *P* < 0.001; N.S, *P* > 0.05 of the Wilcoxon rank sum test).

We next performed subclustering analysis of monocytes and macrophages (n = 27,664 cells) which identified 5 distinct subclusters and confirmed the unique enrichment of M2-like macrophages C1 in the LUAD tissues (**Fig. 4D**, **Supplementary Fig. S18**). Further, we found that C1 M2-like macrophages showed significantly diminished (*P* < 0.001) antigen presentation scores compared to C5 M2-like macrophages which were mainly enriched in normal samples (**Fig. 4D**, **Supplementary Fig. S17B; Supplementary Table S17**). We found markedly reduced expression levels of antigen presentation signature genes with increasing spatial proximity to the LUADs (**Fig. 4E-F**, **Supplementary Fig. S17C**). The spatial pattern of antigen presentation depletion was evident across M2-like macrophages combined from both clusters C1 and C5 and was statistically significant in 4 of the 5 LUAD patients (**Fig. 4G**; *P* < 0.001).

We examined gene and signature score differences between different subsets of DCs (n = 8,694). Spatial patterns were evident in cDC2 and pDC subsets (**Fig. 4H**). We observed differential expression of an inflammatory gene signature between the three cDC2 subclusters with C1, cells of which exhibited the lowest inflammatory scores, heavily enriched in the LUADs (**Fig. 4I-J**; **Supplementary Table S18**). Unsupervised subclustering of cDC2 cells identified three distinct cDC2 subsets characterized by differential expression of an inflammatory signature (highly enriched in C2) and MHC class II genes (enriched in C0/C1) previously shown to discriminate inflammatory from non-inflammatory cDCs ((20); **Supplementary Fig. S17D**). Reduced expression of pro-inflammatory genes and increased levels of anti-inflammatory features were evident among cDC2 cells and along the continuum of normal-to-LUAD space (**Fig. 4K**). cDC2 subcluster with the highest inflammatory score (C2) was further characterized by an inflammatory signature score that was spatially and significantly attenuated with increasing tumor proximity (*P* < 0.001; **Fig. 4L**), and the fraction of C2 among cDC2 subset was overall under-represented in the LUADs (*P* < 0.05; **Fig. 4M**). Notably, the inflammatory signature score significantly and progressively decreased along the continuum from normal lung tissues, to matched premalignant AAHs and invasive LUADs (*P* < 0.01; **Fig. 4N**) in sharp contrast to non-inflammatory DC expression components (**Supplementary Fig. S17E**). We also studied pDC subsets and found spatial enrichment of *FOS, FOSB,* and *JUN*, genes involved in regulating DC immunogenicity (21), with increasing proximity to the tumors (**Supplementary Fig. S17F-G**). We found that a relatively higher score of non-inflammatory to inflammatory expression in cDC2 was associated with prolonged OS and PFI in TCGA LUAD cohort (*P* = 0.009 and *P* = 0.04, respectively) with similar trends observed using an in-house set (MDACC; *P* = 0.2 for OS; **Supplementary Fig. S19**). Altogether, these data underscore spatial immune remodeling in early-stage LUAD that comprises loss of antigen presentation by macrophage subsets and inflammatory phenotypes by DCs. Stromal cells have been shown to impact diverse aspects of tumor immune microenvironment and cancer progression (11,22). Intrigued by the enrichment of stromal subsets in the LUADs (**Fig. 1H**), we further interrogated multiple stromal populations in our dataset (**Supplementary Fig. S20A-C**), including tumor enrichment of vascular smooth muscle cells, adventitial fibroblasts C5, and endothelial cell (EC) venule clusters (**Supplementary Fig. S20D-E**). We pinpointed significantly differentially expressed gene sets in the EC venule subpopulation and altered stromal signatures (**Supplementary Fig. S20F**) that support the observed immune-related changes in the LUAD space, and that are in line with previous observations (11). These comprised tumor-specific activation of extracellular matrix reorganization, syndecan-2 pathway, and neutrophil degranulation, as well as decreased JAK-STAT signaling and reduced antigen-processing and cross-presentation (**Supplementary Fig. S20G-H**).

### Ligand-receptor cell-cell crosstalk in early-stage LUAD

Crosstalk between tumor cells and elements in the TME is implicated in tumor progression largely in part by mediating immunosuppressive phenotypes (23). We utilized iTALK (24) to leverage signals from our scRNA-seq dataset and visualize ligand-receptor (L-R)-mediated intercellular crosstalk (**Fig. 5A**; **Supplementary Table S19**). Computational analysis and annotation were carried out using iTALK’s built-in database focusing on immune checkpoint-receptor pairs (n = 55) and cytokine-receptor pairs (n = 327) (**Fig. 5A**). Overall, we found reduced overlap of L-R interactions between the tumor and distant normal tissues compared to that between the tumor and more proximal (adjacent, intermediate) regions (**Fig. 5B**). We identified altered cellular interactions that were significantly and differentially modulated in LUADs versus their respective spatial normal tissues (**Fig. 5C-F**; **Supplementary Fig. S21A and C**). Among cytokine-receptor pairs, LUADs showed increased communication between *CX3CL1+* tumor epithelial cells and DCs or macrophages expressing increased levels of its cognate receptor *CX3CR1* (**Supplementary Fig. S21A and C; Supplementary Table S20**). *CX3CR1* was increasingly expressed on macrophages and DCs but decreased on CD8 T cells from LUADs (**Supplementary Fig. S21B and D**), in line with previous reports demonstrating pro-tumor features for CX3CR1+ macrophages (25).

**Figure 5.**
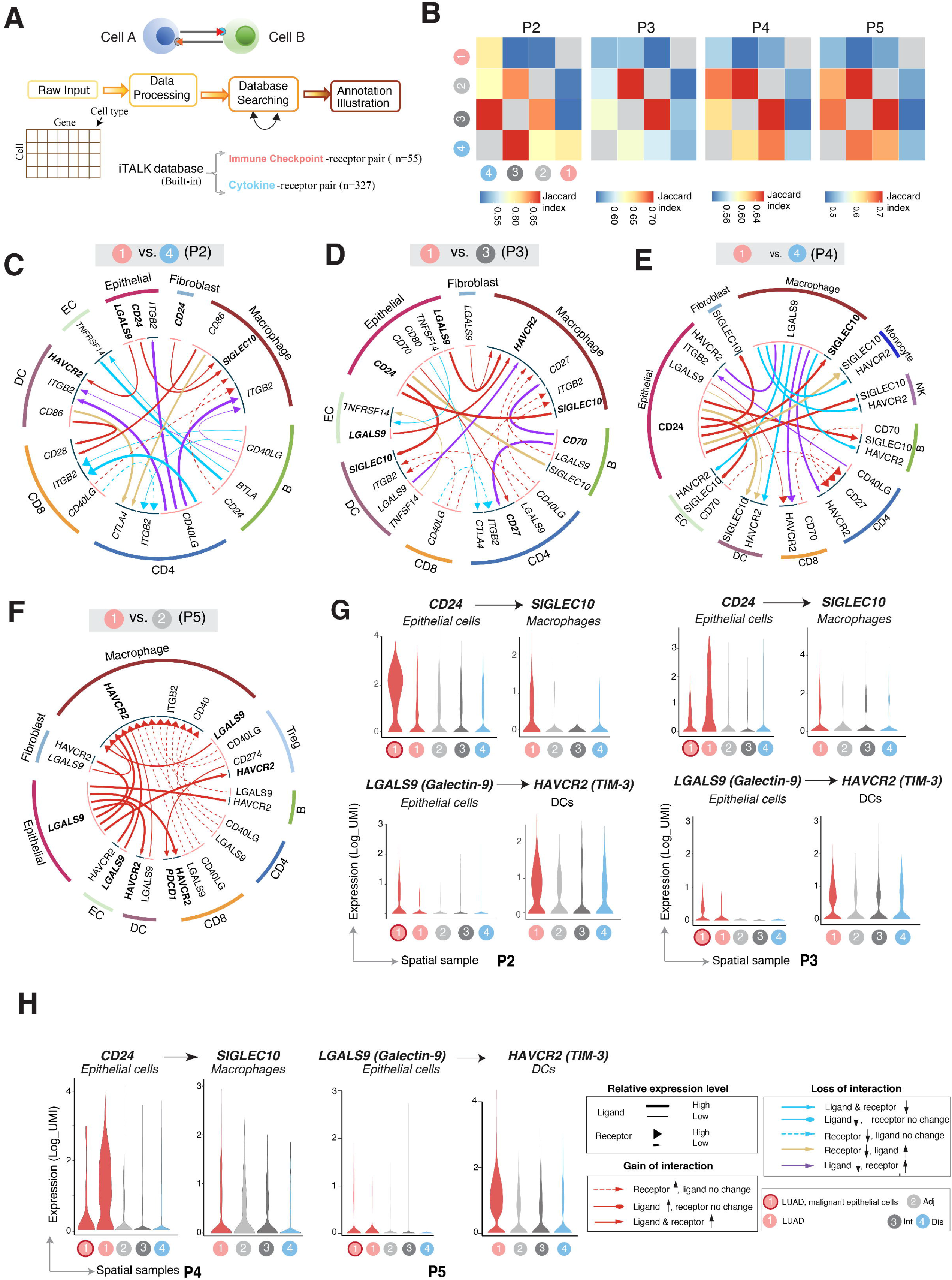
Enriched ligand-receptor cell-cell communication networks between LUADs and their immune microenvironment. **A**, Computational analysis workflow of cell-cell communication using iTALK to identify, from a database of curated ligand-receptor (L-R) pairs, the highly expressed immune checkpoint- and cytokine-receptor pairs, that are significantly and differentially altered (i.e., interactions lost or gained) between LUADs and spatial normal lung tissues. **B**, Heatmaps showing the overlap (quantified by Jaccard index) of predicted ligand-receptor based interactions among individual LUADs and their corresponding spatially distributed normal lung tissues. **C-F**, Representative circos plots showing details of immune checkpoint-mediated L-R pairs compared between each of the LUADs of patients 2 (C), 3 (D), 4 (E) and 5 (F), and selected matching spatial normal lung samples. **G-H**, Violin plots showing expression of the ligand and receptor genes (selected from panels C-F) involving immune checkpoints and showing spatial gain-of-interaction patterns as highlighted in circus plots for patient 2 (panel G left), 3 (panel G right), 4 (panel H left) and 5 (panel H right).

Notably, we identified increased interactions between immune checkpoint proteins *CD24* or *LGALS9 (Galectin-9)* on tumor epithelial cells, and *SIGLEC10* on macrophages or *HAVCR2 (TIM-3)* on DCs, respectively, and which were shared across multiple patients (**Fig. 5C-F**; **Supplementary Table S20**). These interactions were differentially enriched in tumors versus normal tissues at different distances from the LUADs (**Fig. 5G-H**; **Supplementary Fig. S6A**). *CD24* and *LGALS9* expression levels were overall increased in epithelial cells from LUAD tissues, particularly in cells of the malignant-enriched cluster (**Fig. 5G-H**, **Supplementary Fig. S22A**). This pattern of enrichment of *CD24* in cells of the malignant-enriched cluster was also observed in the majority of patients analyzed (**Supplementary Fig. S22B**). Notably, *CD24* expression was also markedly evident in B cells (**Supplementary Fig. S22A-B**) in line with previous reports (26). These findings demonstrate that the early-stage LUAD ecosystem harbors cell-cell communication that confers increased pro-tumor inflammatory and immunosuppressive states.

Our findings above prompted us to validate *CD24* expression patterns using external cohorts. We found progressively and markedly increased expression of the antigen across normal lung tissues, AAHs, and LUADs (**Fig. 6A**; *P* < 0.05). *CD24* positively correlated with expression of the epithelial marker *EPCAM* as well as with levels of pro-tumor and immune suppressive features (*TIGIT, CTLA4, FOXP3, CCL19*), in contrast to negatively correlating with anti-tumor immune markers (*GZMB, GZMH, PRF1*) (**Fig. 6B**, **Supplementary Fig. S23A**). These observations were, for the most part, further validated in LUADs and matched normal tissues from TCGA cohort (27) (**Fig. 6C-D**, **Supplementary Fig. S23B**). Using an in-house cohort of early-stage LUADs analyzed by targeted immune profiling (see Supplementary Methods), we found that relatively higher *CD24* was associated with shortened OS and PFI (**Fig. 6E**; *P* = 0.07). Consistent with the above, *CD24* expression positively correlated with that of *EPCAM* (*r* = 0.31) and negatively correlated with both *PRF1* (*r* = −0.37) and immune cytotoxicity score ((28), *r* = −0.33; **Fig. 6F**). Further, analysis of an independent in-house tissue microarray of treatment-naïve LUADs revealed that CD24 immunohistochemical protein expression was prevalent in tumor cells (**Fig. 6G** left) and, in line with the above, was also associated with reduced OS (*P* = 0.04) and PFI (*P* = 0.007) (**Fig. 6G** right). We further found in a syngeneic mouse model that loss of *Cd24a* by CRISPR mediated knockout or its inhibition using neutralizing antibodies significantly reduced the growth of mouse LUAD cells *in vivo* (Supplementary Methods**, Supplementary Fig. S24, Fig. 6H**). These findings show that CD24, which we found to be at the core of an enriched cell-cell interactome in early-stage LUAD, is associated with a pro-tumor immune contexture and poor prognosis as well as promotes LUAD growth *in vivo*.

**Figure 6.**
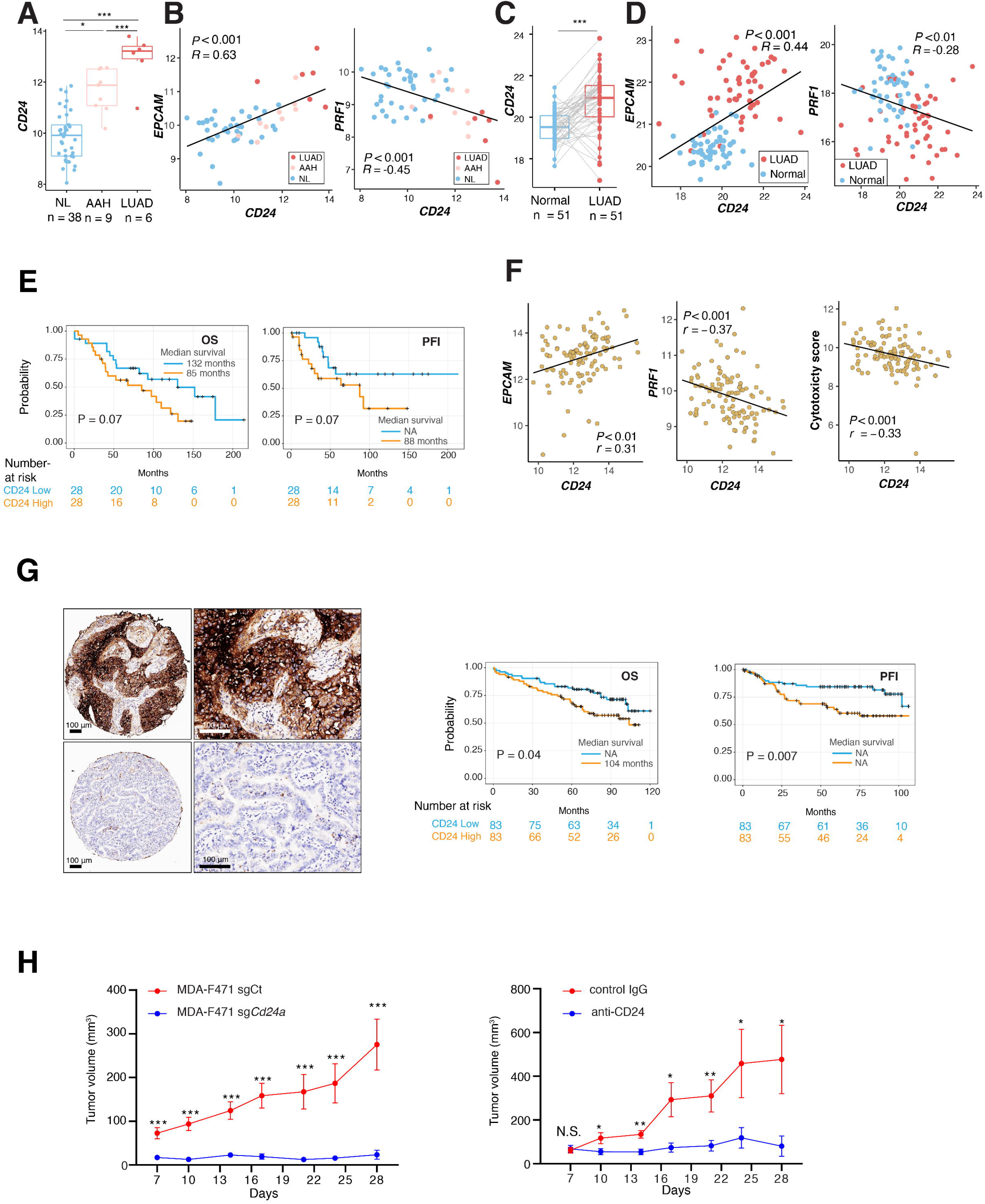
Pro-tumor phenotypes associated with augmented CD24 in LUAD. **A,** Boxplot showing *CD24* expression levels in an independent cohort of normal lung tissues (NL), premalignant atypical adenomatous hyperplasias (AAH) and LUADs assessed using the Nanostring immune Counter panel (see Supplementary Methods) (*, *P* < 0.05; **, *P* < 0.01; ***, *P* < 0.001 of the Wilcoxon rank sum test). **B**, Scatterplots using Pearson correlation coefficients between levels of *CD24* with *EPCAM* and *PRF1* in the NL, AAH, and LUAD samples. **C**, Boxplot depicting *CD24* expression levels in LUADs and matched normal lung (NL) tissues from TCGA LUAD cohort (***, *P* < 0.001 of the Wilcoxon rank sum test). **D**, Scatter plots showing correlation of expression using Pearson’s correlation coefficients between *CD24* and *EPCAM* or *PRF1*. **E,** Overall survival (OS) and progression-free interval (PFI) in a subset of early-stage LUAD patients analyzed by targeted immune profiling (MDACC cohort) and dichotomized by median *CD24* mRNA expression (*CD24* low n = 28 and high n= 28). Survival analysis was performed using Kaplan–Meier estimates and two-sided log-rank tests. **F,** Scatter plots showing correlation of expression using Pearson’s correlation coefficients between *CD24* and *EPCAM*, *PRF1* or cytotoxicity score in MDACC cohort. Cytotoxicity score was calculated as the square root of the product of *GZMB* and *PRF1* expression. **G,** Representative images showing relatively high (top) and low (bottom) immunohistochemical CD24 staining (left). Scale bar = 100 µm. OS (middle) and PFI (right) analysis in the early-stage LUAD TMA. Patients were dichotomized based on median CD24 protein expression (CD24 low n = 83 and high n = 83). The analysis was performed with the Kaplan–Meier estimates and two-sided log-rank tests. **H**, *In vivo* growth of MDA-F471 cells subcutaneously implanted into syngeneic mice. Lengths and widths of tumors were measured twice per week for 3 weeks and tumor volumes were calculated according to the formula (length x width^2^)/2, and tumor growth was plotted as mean ± SEM. Mice in the left panel were implanted with MDA-F471 sgCt or MDA-F471 sg*Cd24a* cells following FACS-sorting for high or low expression of CD24 surface protein, respectively. Mice in the right panel were implanted with parental MDA-F471 cells and treated with either control IgG or anti-CD24 antibody at the indicated timepoints (*, *P* < 0.05; **, *P* < 0.01; ***, *P* < 0.001; unpaired Student’s *t* test).

## DISCUSSION

Molecular changes have been documented in the local niche of LUAD including loss-of-heterozygosity in 3p and 9p, point mutations and tumor suppressor methylation (29,30). Earlier work also underscored transcriptome profiles, somatic driver variants, as well as genome-wide allelic imbalance events that are shared between lung cancer and adjacent normal-appearing airway cells but that are absent in distant normal cells, thereby pointing to putative drivers of lung oncogenesis (4,5). These earlier studies have focused on understanding geospatial profiles by bulk profiling methods, thereby inadvertently obscuring the individual contributions of epithelial and TME cues to lung cancer pathogenesis. Our knowledge of the geospatial architecture of individual cell populations in early-stage LUAD evolution remains poorly understood. By single-cell interrogation of a unique multi-region sampling design with epithelial cell enrichment, we here characterized spatial and ecological maps comprising various epithelial and non-epithelial subsets that underlie emergence of early-stage LUAD from its local niche.

Multi-regional or spatial analyses have been employed to interrogate intratumor heterogeneity (ITH) in solid tumors, including LUADs, in order to understand evolutionary trajectories and therapy response (31). Our analyses showed that ITH is evident at both the tumor epithelial cell and intra-site levels, i.e., within the same tumor region. We applied an integrative approach to dissect ITH of likely malignant cells and characterized cell clusters with differential transcriptomic profiles, evolutionary trajectories, CNV burdens, and/or driver mutations. We also found “normal” cells in the LUAD tissues themselves that are close in the inferred trajectory paths to specific malignant-enriched subsets, possibly representing tumor cells-of-origin. Normal basal cells (expressed *KRT5, KRT15, KRT17,* and *P63*) were consistently evident in the LUAD samples, possibly corroborating previously reported basal differentiation hierarchies in the normal lung (32,33). It is noteworthy that we found, in the normal-appearing samples, cells with features of malignant-enriched subsets and heterogeneous CNV profiles. Whether these cells comprise early LUAD precursors, mutagenic clones that do not progress to malignancy, or putative molecular field cancerization remains to be investigated (34).

LUADs exhibit remarkable inter-patient heterogeneity in histological differentiation patterns, driver alterations and inferred tumor cells-of-origin (35). In our cohort, we identified malignant gene expression features that varied between the patients and that pointed to likely distinct tumor cells-of-origin and/or LUAD histopathological patterns. We pinpointed G12D mutations in *KRAS* to a unique subset of the malignant-enriched cluster within P2 LUAD. These cells had overall low CNVs, perhaps reminiscent of findings in LUADs driven by strong driver genes *in vivo* (36). These cells exhibited increased expression of genes associated with *KRAS*-mutant cancer such as *LCN2* (a marker of inflammation, (17)), and reduced expression of the lineage-specific oncogene *NKX2-1* (18), in line with expression patterns in mucinous *KRAS*-mutant LUADs (e.g., P2). Compared to other C9 (malignant-enriched cluster) cells from the same LUAD (P2), *KRAS*-mutant cells showed reduced expression of airway lineage-specific genes (e.g., *SCGB3A1*, *SFTPB*) in accordance with recent reports (8). Our findings also allude to the possibility that tumor cell lineage plasticity may occur at an early-stage of *KRAS*-mutant LUAD carcinogenesis – a supposition that warrants exploration of a larger and more diverse repertoire of *KRAS*-mutant cells. Of note, we could not characterize the full spectrum of ITH, clonality, and inter-patient heterogeneity given the limited number of patients profiled and the use of scRNA-seq which is not ideal for examining tumor clonality or genomic alterations. Nonetheless, our in-depth analysis of a relatively large number of epithelial cells unveiled different characteristics (airway lineages, malignant programs, potential tumor cells-of-origin, and cellular ITH) of the epithelial architecture of early-stage LUAD. These insights could be further extended to profile a larger and more diverse array of LUADs using anticipated advances enabling simultaneous scRNA-seq and scDNA-seq of the same cell.

Earlier studies have shown that immunosuppressive T regulatory cells are crucial for immune evasion in lung cancer (37,38). We found Tregs co-expressing both *TIGIT* and *CTLA-4* immune checkpoints and that were progressively enriched with increasing geospatial proximity to the LUADs – suggesting a value in combinatorial targeting of multiple checkpoints for immunotherapy of early-stage LUAD (39). scRNA-seq analysis of our limited patient cohort also pinpointed important attributes to other lymphoid populations. We identified B cell signatures that are spatially enriched in the LUADs, progressively increased along the course of normal to preneoplasia and invasive LUAD and associated with prolonged survival. These data suggest important yet unexplored roles for B cell phenotypes in immune evolution of LUAD (40). We also noted strikingly high plasma cell fractions among LUADs from smokers, suggesting that plasma cells may play important roles in the pathogenesis of smoking-associated LUAD and its microenvironment (41). Our analysis also pointed to mechanisms by which the myeloid immune microenvironment permits LUAD pathogenesis. Tumor-specific M2-like macrophages displayed diminished antigen presentation, whereby expression levels of MHC genes as well as genes involved in peptide transport and loading (*TAP1, TAP2, TAPBP;* (42)) were markedly reduced. We also found DC subclusters and signatures that were recently reported in healthy donors and in cancer (20,43). These included a shift from an inflammatory to a non-inflammatory DC state across normal lung to premalignant AAH up to LUAD and that was associated with prolonged survival in patients -- thus pointing to immune cues that could perhaps be harnessed to manipulate the immunogenicity of tumors (43). We also identified an inflammatory tumor-depleted cDC cluster with increased expression of scavengers such as *CD36*. Dendritic CD36 permits acquisition and presentation of cell surface antigens (44), and the precise effects of its stark absence in a LUAD-specific cDC subset the tumor immune microenvironment warrant further investigation. Nevertheless, these findings suggest that unique cDC subsets play critical roles in LUAD pathogenesis and, thus, could be potential targets for immune-based interception.

Deciphering ligand-receptor mediated interactions can elucidate cell-to-cell communication implicated during carcinogenesis, tumor-immune co-evolution and immune reprogramming, and thus, may help identify potential immune therapeutic targets (23). In this study, we applied the iTALK tool (24) developed by our group and performed a deep analysis of cellular interaction networks. We identified significant immune checkpoint- (e.g., *CD24*, *Galectin-9*, or *TIM-3)* and cytokine- (e.g., *CX3CL1)* receptor interactions whose enrichment or depletion in the LUAD space signified a highly pro-tumorigenic milieu. These findings are in accordance with the CD24–Siglec-10 interaction and subsequent “do not eat me” signal recently highlighted in breast cancer (26). Additionally, our findings on CD24 including prominent expression in tumor epithelium, association with pro-tumor immune phenotypes and reduced survival, and functional role in LUAD *in vivo*, suggest that CD24 may be a viable target for treatment of early-stage LUAD. Overall, our interactome analyses implicate potential targets as culprits in LUAD pathogenesis.

In summary, our results provide a spatial atlas of early-stage LUAD and its nearby and distant lung ecosystem. This atlas comprises high cellular heterogeneity as well as spatial dynamics in cell populations, cell states and their transcriptomic features that underlie LUAD evolution from the peripheral lung ecosystem. Our extensive transcriptomic dataset of lung epithelial and immune cells, other populations such as stromal and endothelial subsets, as well as of tumor-pertinent cell-cell interactions, constitutes a valuable resource to functionally interrogate LUAD trajectories at high resolution and generate strategies for its early treatment. Also, our study’s multi-region sampling design in conjunction with single-cell analysis could help address specific questions in early malignant and immune biology of other solid tumors.

## METHODS

Additional description of methods can be found in the Supplementary Data file.

### Multi-region sampling of surgically resected LUADs and normal lung tissues

Patients undergoing surgical resection for primary early-stage lung adenocarcinoma (I-IIIA) were carefully selected for derivation of multi-region samples for single cell analysis (**Supplementary Table S1**). All patients were evaluated at the University of Texas MD Anderson Cancer Center and had provided informed consents under approved institutional review board protocols. Immediately following surgery, resected tissues were processed by an experienced pathologist assistant (PB). One side of the specimen was documented and measured, followed by tumor margin identification. Based on the placement of the tumor within the specimen, incisions were made at defined collection sites in one direction along the length of the specimen and spanning the entire lobe: tumor-adjacent and -distant normal parenchyma at 0.5 cm from the tumor edge and from the periphery of the overall specimen/lobe, respectively. An additional tumor-intermediate normal tissue was selected for P2-5 that ranged between 3-5 cm from the edge of the tumor. Sample collection was initiated at normal lung tissues that are farthest from the tumor moving inwards towards the tumor to minimize cross-contamination during collection.

### scRNA-seq analysis

Tumor and spatial normal parenchyma tissues (n = 19 samples) were immediately transported on ice for mincing and enzymatic digestion (Supplementary Methods). Cells were sorted (by FACS) for viable singlets cells (and also EPCAM+/− fractions from P2-P5), followed by processing for scRNA-seq library construction using 10X Genomics and sequencing using Novaseq6000 platforms. Single-cell analyses were performed using available computational framework. Raw scRNA-seq data were pre-processed, demultiplexed, aligned to human GRCh38 reference and feature-barcodes generated using CellRanger (10X Genomics, version 3.0.2). Details of quality control including quality check, data filtering, as well as identification and removal of cellular debris, doublets and multiplets are found in Supplementary Methods. Following quality filtering, a total of 186,916 cells were retained for downstream analysis. Raw unique molecular identifier (UMI) counts were log normalized and used for principal component analysis using Seurat (45). Batch effects were statistically assessed using k-BET (46) and corrected by Harmony (47) (Supplementary Methods), followed by unsupervised clustering analysis using Seurat v3 (45). Uniform Manifold Approximation and Projection (UMAP) (48) was used for visualization of clusters. Clustering robustness was examined for major clusters and subclusters (see Supplementary Methods), and all subclustering analyses were performed separately for each compartment. For unsupervised clustering analysis, we applied single-cell consensus clustering (SC3; (49)) approach using default parameters and independently of cell lineage annotation. Differentially expressed genes (DEGs) for each cell cluster were identified using the *FindAllMarkers* function in Seurat R package. We defined cell types and cluster functional states by integrating the enrichment of canonical marker genes, top-ranked DEGs in each cell cluster, and the global cluster distribution.

To study hierarchical relationships among cell types, pairwise Spearman correlations were calculated from average expression levels (Seurat function *AverageExpression*), based on which Euclidean distances were calculated. Monocle 2 (version 2.10.1; (50)) was applied to construct trajectories. Likely malignant cells were distinguished from non-malignant subsets based on information integrated from multiple sources including cluster distribution of the cells, gene expression, as well as presence of mutations (*KRAS* in P2, see Supplementary Methods) and CNVs. CNVs were inferred using inferCNV tool (13) with NK cells as the control. To evaluate the robustness of CNV inference, T and B cells were also used as controls. To identify significant ligand-receptor pairs among major cell lineages, the top 30% of most highly expressed genes were included in the analysis. Significant cellular interactions were identified using iTALK as previously described (24). For ligand-receptor annotation, the iTALK built-in ligand-receptor database was used (https://github.com/Coolgenome/iTALK).

### Statistical analysis

All statistical analyses were performed using R package v3.6.0. Pseudo-bulk gene expression values for defined cell clusters were calculated by taking mean expression of each gene across all cells in a specific cluster. Pearson’s correlation analysis was used to identify genes significantly correlated with *CD24* expression. Log-rank tests and Kaplan-Meier plots were used for survival analysis. All statistical significance testing was two-sided, and results were considered statistically significant at *P* < 0.05. The Benjamini-Hochberg method was applied to control the false discovery rate (FDR) in multiple comparisons and to calculate adjusted *P –* value (q-values).

### Data Availability

All sequencing data generated in this study are deposited in the European Genome-phenome Archive (EGAS00001005021).

