## Supplementary Methods for "Resolving the spatial and cellular architecture of lung adenocarcinoma by multi-region single-cell sequencing"

**SUPPLEMENTARY DATA**

**Supplementary Methods**

**Single cell isolation from LUADs and normal spatial samples**

Fresh tissues were collected in RPMI medium supplemented with 2% fetal bovine serum (FBS) and maintained on ice for immediate processing. Tissues were placed in a cell culture dish containing Hank’s balanced salt solution (HBSS) on ice and extra-pulmonary airways and connective tissue were removed with scissors. Samples were transferred to a new dish on ice and minced into tiny pieces (approximately 1mm^3^) followed by enzymatic digestion using a cocktail containing Collagenase A (10103578001, Sigma Aldrich), Collagenase IV (NC9836075, Thermo Fisher Scientific), DNase I, (11284932001, Sigma Aldrich), Dispase II (4942078001, Sigma Aldrich), Elastase 1 (NC9301601, Thermo Fisher Scientific) and Pronase (10165921001, Sigma Aldrich), as previously described (1). Samples were incubated in a 37 °C oven for 10 minutes with gentle rotation, pipet mixed at room temperature for further disaggregation, and then incubated at 37 °C for an additional 10 minutes. Samples were then filtered through 70 μm strainers (Miltenyi biotech, 130-098-462) and washed with ice-cold HBSS. Filtrates were then centrifuged and re-suspended in ice-cold ACK lysis buffer (A1049201, Thermo Fisher Scientific) for red blood cell (RBC) lysis. Following RBC lysis, samples were centrifuged and re-suspended in ice-cold FBS, filtered (using 40 μm FlowMi tip filters; H13680-0040, Millipore), and an aliquot was taken to enumerate cells and check for viability by Trypan blue exclusion analysis.

**Sorting and enrichment of viable lung EPCAM+ epithelial and EPCAM- non-epithelial singlets**

Single cells from patient 1 (P1) were stained with SYTOX Blue viability dye (S34857, Life Technologies), and processed on a fluorescence-activated cell sorting (FACS) Aria I instrument. Cells from patients 2 through 5 (P2-P5) were stained with anti-EPCAM-PE (347198, BD Biosciences;1:50 dilution in ice-cold phosphate-buffered saline, PBS, containing 2% FBS) for 30 minutes with gentle rotation at 4°C. EPCAM-stained cells were then washed, filtered using 40 μm tip filters, stained with SYTOX Blue and processed on a FACS Aria II instrument. Doublets and dead cells were eliminated, and two populations of viable (SYTOX-negative) singlets were separately collected in PBS containing 2% FBS: EPCAM-positive (EPCAM+, epithelial) and EPCAM-negative (EPCAM-, non-epithelial). Cells were washed again to eliminate ambient RNA, and a sample was taken for counting by Trypan Blue exclusion (T8154, Sigma Aldrich) before loading on 10X Chromium microfluidic chips.

**Preparation of single-cell 5’ gene expression libraries**

Up to 10,000 cells per sample were partitioned into nanoliter-scale Gel beads-in-emulsion (GEMs) using Chromium Next GEM Single Cell 5' Gel Bead Kit v1.1 (1000169, 10X Genomics) and by loading onto Chromium Next GEM Chips G (1000127, 10X Genomics). GEMs were then recovered to construct single-cell 5’ gene expression libraries using the Chromium Next GEM Single Cell 5’ Library kit (1000166, 10X Genomics) according to the manufacturer’s protocol. Briefly, recovered barcoded GEMs were broken and pooled, followed by magnetic bead clean-up (Dynabeads MyOne Silane, 37002D, Thermo Fisher Scientific). 10X-barcoded full-length cDNA was then amplified by PCR and analyzed using Bioanalyzer High Sensitivity DNA kit (5067-4626, Agilent). Up to 50 ng of cDNA was carried over to construct gene expression libraries and was enzymatically fragmented and size-selected to optimize the cDNA amplicon size prior to 5’ gene expression library construction. Further, samples were subject to end-repair, A-tailing, adaptor ligation, and sample index PCR using Single Index Kit T Set A (2000240, 10X Genomics) to generate Illumina-ready barcoded gene expression libraries. Library quality and yield was measured using Bioanalyzer High Sensitivity DNA (5067-4626, Agilent) and Qubit dsDNA High Sensitivity Assay (Q32854, Thermo Fisher Scientific) kits. Indexed libraries were normalized by adjusting for the ratio of the targeted cells per library as well as individual library concentration and then pooled to a final concentration of 10 nM. Library pools were then denatured and diluted as recommended for sequencing on the Illumina NovaSeq 6000 platform.

**Raw single-cell sequencing data processing and quality control**

Detailed quality control metrics were generated and carefully evaluated. Cells with low complexity libraries or likely cellular debris (in which detected transcripts are aligned to less than 200 genes) were filtered out. Low-quality cells where >15% of transcripts were derived from the mitochondrial genome were also excluded (**Supplementary Fig. S2A** middle panel). Doublets were identified for exclusion using a multi-step approach: 1) library complexity: cells with highly complex libraries (in which detected transcripts are aligned to more than 6500 genes); 2) cluster distribution: doublets or multiplets likely forming distinct clusters with hybrid expression features and exhibiting an aberrantly high gene count; 3) cluster marker gene expression: cells of a cluster expressing markers from distinct lineages (e.g., cells in the T-cell cluster showing expression of epithelial cell markers). Step 3 was performed after batch effect assessment and correction (see below). The above steps were repeated multiple times to ensure elimination of most barcodes associated with cell doublets.

We then used k-BET (2) for batch effect assessment and applied Harmony (3) for actual batch effect correction. As outlined below, these tools were applied to clustering of major lineages and specific subsets before any clustering analysis or cell type identification/annotation was performed. Statistical assessment of possible batch effects was performed using the R package k-BET (a robust and sensitive k-nearest neighbor batch-effect test) (2). k-BET was run on cells from all patient samples including P1, and on major lymphoid cell types including B cells, CD4 T cells, CD8 T cells, myeloid cells, and epithelial cells separately with default parameters. Each cell type was down sampled to 500 cells. We chose the k input value from 1% to 100% of the sample size. In each run, the number of tested neighborhoods was 10% of the sample size. The mean and maximal rejection rates were then calculated based on a total of 100 repeated k-BET runs. A low rejection rate indicates homogeneous mixing of samples from different batches. Following estimation of sample processing- or sequencing-related batch effects using k-BET, and to remove batch effects, we next employed Harmony (3) before clustering analysis (**Supplementary Fig. S2A, Fig. S5A, Fig. S11B, Fig. S16B, Fig. S20A**).

**Determination of major cell types and states as well as inference of cell cycle stage**

Cell type identification was performed on Harmony-defined clusters following batch effect correction. To determine the major cell type of each cell, differentially expressed genes (DEGs) were identified for each cell cluster using the *FindAllMarkers* function in Seurat and the top 50 most significant DEGs were reviewed. In parallel, feature plots were generated for top 50 DEGs and for a suggested set of canonical immune and stromal cell markers, as described in previous reports (4-7), followed by a manual review process. Enrichment of these markers (e.g., *EPCAM* for epithelial cells; *PTPRC* for immune cells; *CD3D/E* for T cells; *CD8A/B* for CD8 T cells, *CD4/CD40LG* for CD4 T cells; *CD19/MS4A1/CD79A* for B cells, *COL1A1/COL1A2* for fibroblasts) was considered a strong indication of the clusters representing the corresponding cell types. Top-ranked DEGs and the enrichment of canonical marker genes were integrated with global cluster distribution to infer major cell types and cellular states, as previously described (8,9). Cell cycle stage was computationally assigned for each individual cell by the function *CellCycleScoring* that is implemented in Seurat. Cell cycle stage was inferred based on expression profiles of the cell cycle related signature genes, as previously described (10). For transcriptional signature analysis, single-sample GSVA (ssGSVA) was applied to the scRNA-seq data and pathway scores were calculated for each cell using *gsva* function in GSVA software package (11).

**Analysis of clustering robustness**

Multiple methods were applied to assess the robustness of the clustering results in this study. In addition to Harmony, reciprocal PCA (rPCA) (12) was used as an independent integration method to cluster and subcluster multiple cellular subsets. Clustering consistency was quantified by calculating the Jaccard index as a measure of the overlap between the results obtained with Harmony and those obtained with rPCA. Due to the large number of cells in the analysis of the five major lineages (n = 186,916 cells in total), we randomly sampled 75% of all cells in rPCA analysis for computational efficiency (**Supplementary Fig. S3A**). To evaluate the extent with which the clustering results relied on individual cells being included, we analyzed randomly down-sampled cells: 25%, 50% and 75% from the original dataset for analysis of major lineages, EPCAM+ epithelial cells, CD8+ T cell subsets, and monocytes/macrophages. Results from clustering of down-sampled cells were compared with those derived from all cells using Jaccard index (**Supplementary Fig. S3B, Fig. S5C, Fig. S12, Fig. S18**).

**Analysis of large-scale copy number variations and *KRAS* mutation status**

To quantify the level of aneuploidy, profiles of copy number variation (CNV) generated by inferCNV (<https://github.com/broadinstitute/inferCNV> and as described in Methods section) were aggregated using a strategy similar to that described in a previous study (13). We first computed arm-level CNV scores as the mean of the squares of CNV values across each chromosomal arm. The arm-level CNV scores were further aggregated across all chromosomal arms by taking the arithmetic mean value of the arm-level scores using R function *mean*. To associate CNV clusters with tumor clonal architecture, we constructed phylogenetic neighbor-joining trees using the R package phangorn for cells from P3 LUAD (14). The consistency between the phylogenetic neighbor-joining tree and the original hierarchical clustering tree was quantified by calculating the Pearson’s correlation coefficient between the orders of cells (clades and clusters) in both trees. *P* - values were calculated using *cor.test* function.

For single-cell somatic mutation analysis, we were able to specifically capture and analyze *KRAS* codon 12 mutation status in our cohort since *KRAS* codon 12 falls within the genomic bandwidth of our 5’ single-cell gene expression assay, i.e., its locus is present within the first 100 bp from the 5’ end of transcripts sequenced. Reads from P2 samples were extracted from the original BAM files using cell-specific barcodes and genomic coordinates of *KRAS* codon 12 mutations and then subjected to quality filtering and duplicates removal, mutation identification and annotation. Extracted alignments were manually evaluated using IGV (15).

**Analysis of LUAD expression datasets**

Gene signatures (B cell C0 signature, non-inflammatory versus inflammatory DC signature from Villani et al (8)) were validated in a previously published dataset by our group comprised of normal lung tissues, precursor atypical adenomatous hyperplasias (AAHs) and matched LUADs (n = 15, each) (16). Gene expression matrix with log transformed TPM values was downloaded from GSE102511. Pearson’s correlation coefficients between different genes were calculated using log2 transformed TPM values.

Correlation between expression of *CD24* and that of other immune-related genes or *EPCAM* was validated in three independent datasets. Firstly, we analyzed select genes of interest in a cohort of premalignant and invasive lung lesions (n = 9 AAHs, n = 6 LUADs) and normal lung tissues (n = 38) that were profiled using the nCounter PanCancer Immune Profiling Panel (NanoString Technologies) (17). Secondly, we interrogated bulk RNA-seq expression data of 51 treatment-naïve early-stage LUADs (disease stages: I, II and III) LUADs with matched adjacent normal lung tissues from The Cancer Genome Atlas (TCGA) cohort (18) and downloaded from the NCI Cancer Genomic Data Commons (NCI-GDC: <https://gdc.cancer.gov)>. For these two datasets, Pearson’s correlation coefficients were computed using normalized expression data (log2 transformed FPKM values). Thirdly, expression of select genes was validated in an in-house LUAD cohort (MDACC; n = 109) analyzed by targeted immune profiling of 1,392 genes using the HTG EdgeSeq Precision Immuno-Oncology Panel (HTG Molecular Diagnostics, Inc.). The R package MCP-counter (19) was applied to the normalized data of primary LUADs in both TCGA and MDACC cohorts. For MDACC cohort, we also performed correlation analysis of *CD24* with a computed immune cytotoxicity score (20), and Pearson’s correlation coefficient was calculated using median normalized expression.

**Immunohistochemical analysis of CD24 in LUAD**

Archived and surgically resected tumor specimens were collected from treatment-naïve LUAD patients (n = 240) with early-stage (stages I-III) disease and incorporated into a tissue microarray (TMA). Patients in this cohort were evaluated at The University of Texas MD Anderson Cancer Center and consents were obtained from all patients and were approved by the institution’s review board. Protein expression of CD24 was evaluated by immunohistochemistry (IHC) using a Leica Bond Max automated stainer (Leica Biosystems Nussloch GmbH) and using the same conditions as simultaneously stained positive and negative controls. Briefly, 4 μm-thick sections from this TMA were deparaffinized and rehydrated with standard Leica Bond protocol followed by antigen retrieval at 100°C for 20 minutes with Bond ER Solution #1, (Leica Biosystems, equivalent Citrate Buffer, pH6). Anti-CD24 primary antibody (Clone SN3, dilution 1:200, Invitrogen Cat# MA5-11828) was incubated for 15 minutes at room temperature and detected using the Bond Polymer Refine Detection kit (DS9800, Leica Biosystems) with DAB as chromogen. Slides were then counterstained with hematoxylin, dehydrated, and mounted. The stained slides were digitally scanned using the Aperio ScanScope Turbo slide scanner (Leica Microsystems Inc.) and images were captured at ×200 magnification. The images were visualized using ImageScope software (Leica Microsystems, Inc.), and digital image analysis was performed using the Aperio Image Toolbox (Leica Microsystems Inc.) by an experienced pathologist (JF). Cytoplasmic expression of CD24 was quantified using a 4-value intensity score (0, none; 1, weak; 2, moderate; and 3, strong) and the percentage (0%–100%) of the extent of reactivity. A final expression score (H-score) was obtained by multiplying the intensity and reactivity extension values (range, 0–300) as described previously (21,22). TMA specimens were simultaneously stained with positive and negative controls using the same IHC conditions with a Bond Max automated staining system (Leica Microsystems, Inc.).

**Survival analysis**

Survival analysis was performed in treatment-naïve early-stage LUAD patients from TCGA (n = 468) as well as in 56 patients (MDACC cohort) who had not received adjuvant therapy following surgery. Patients were dichotomized into upper (Q1) and lower (Q4) quartiles expressing immune cell signatures analyzed in our single-cell cohort, namely B cell C0 signature score and cDC2 signature score of the expression of a non-inflammatory/inflammatory ratio. For survival analysis based on CD24 expression, patients were dichotomized based on median *CD24* mRNA or protein expression. Statistical differences in survival between different groups were evaluated in R using the log-rank test and the Kaplan–Meier method for estimation of survival probability.

***In vitro* knockdown of *Cd24a* expression in mouse LUAD cells**

Early passage and mycoplasma-free MDA-F471 cells previously derived from LUADs of *Gprc5a^-/-^* mice (23) were maintained in a humidified incubator in DMEM-F12 medium supplemented with 10% fetal bovine serum (FBS) and antibiotics. Knockout of *Cd24a* expression in MDA-F471 cells was established using the CRISPR-Cas9 system, using lentiviral supernatant from predesigned Edit-R Lentiviral small guide RNAs (sgRNAs) for guiding Cas9 nuclease to create double-strand breaks in the target DNA. Cells were transduced with either control non-targeting sgRNA (VSGC11954; sgCt) or a combination of two different target sequences of mouse *Cd24a* (VSGM11945-15EG12484; sg*Cd24a*; both Horizon Discovery, Boyertown, PA) according to manufacturer’s protocol. At 48 hours post-transduction, cells were treated with puromycin (5 µg/ml) for selection of cells with integrated sgRNA, and selection medium was replenished every three days. At day 10 post-transduction, cells with high transduction efficiency were sorted by FACS.

**Fluorescence-activated cell sorting of transduced cells based on CD24 surface protein expression**

sgCt- and sg*Cd24a-*transduced MDA-F471 cells were trypsinized and washed with PBS containing 2% FBS. Cells were stained with mouse-specific Pe-Cy7-conjugated anti-CD24 antibody (1:200 dilution, clone M1/69, eBioscience) for 30 mins in the dark and with gentle rotation at 4°C. Cells were then washed twice, filtered, stained with SYTOX Blue viability dye, and processed on a FACS Aria I instrument. Under sterile conditions, CD24^high^-expressing MDA-F471 were sorted from cells transduced with sgCt, and CD24^low^ cells were sorted from cells transduced with sg*Cd24* targeting lentiviruses. A fraction of the sorted cells was used for quantitative real time PCR analysis, and the rest of the cells were cultured in a humidified incubator and maintained in puromycin selection media.

**Quantitative real time-PCR** (**qRT-PCR**) **analysis of *Cd24a* expression in sorted mouse LUAD cells**

RNA was extracted from sorted MDA-F471 cells using the RNeasy extraction kit (Qiagen) following the manufacturer’s instructions, and the purified RNA was assessed on Nanodrop (RNA quality) and quantified using Qubit RNA Broad Range Assay (Q10211). cDNA was reverse transcribed from total RNA using the High-Capacity cDNA reverse transcription kit (Applied Biosystems, Foster City, CA). *Cd24* was amplified using Taqman probes specific for mouse *Cd24a* (Mm00782538_sH FAM-MGB) and TaqMan Fast Advance Master Mix (4444963, Applied Biosystems) on QuantStudio 7 Flex Real-Time PCR System (Applied Biosystems), and according to the following cycling conditions: 1s denaturation at 95°C, 20s annealing and extension at 60°C for 40 cycles. Gene expression was normalized to both *Gapdh* (Mm99999915_g1 FAM-MGB) and *Actin* (#Mm00607939_s1 FAM-MGB) using the 2^-ΔΔCt^ method.

**Animal housing**

Animal experiments were conducted according to Institutional Animal Care and Use Committee-approved protocols. *Gprc5a^−/−^* mice with a mixed C57BL/6 x 129 genetic background were generated as previously described (24,25) and were maintained in a pathogen-free animal facility. Mice were continually monitored to ensure adequate body condition scores and to ensure that tumors were less than 2 cm^3^ in diameter and that there was less than 50% ulceration. For all experiments, 8-week old age-matched *Gprc5a^−/−^* female mice were divided into starting groups of 7 - 10 mice for each experimental arm.

**Analysis of LUAD transplant growth in syngeneic mice following genetic deletion of *Cd24* or inhibition with neutralizing antibodies**

Before injection, parental MDA-F471, MDA-F471-CD24^hi^ or MDA-F471-CD24^neg^ cells were trypsinized, washed in PBS, and re-suspended at an average density of 1 x 10^6^ cells in 50 µl PBS (calcium- and magnesium-free) supplemented with 50% phenol red-free Matrigel Matrix with high protein concentration (Corning, NY). Female mice received analgesic (Buprenorphine SR, 5 µg/ml, intraperitoneally) and were anesthetized with isoflurane. Under aseptic conditions and using a 3/10-cc insulin syringe equipped with a 30-G hypodermic needle, cells in Matrigel suspension were engrafted into the right flank of each mouse. Subcutaneous tumors were measured twice a week starting at day 7 post-engraftment using digital calipers, and tumor volume was calculated using the formula: (length x width^2^)/2 and normalized to XXX. Mice engrafted with parental MDA-F471 cells were randomized into treatment groups (7-10 mice per group), receiving either anti-CD24 monoclonal antibody or rat IgG1 isotype control (200 µg), starting at day 7 post-engraftment, twice a week for 3 weeks (6 doses in total). At day 28 post-engraftment, mice were sacrificed and subcutaneous plugs were collected for subsequent RNA extraction.

**Supplementary Figure Legends**

**Supplementary Fig. S1. Schematic view of the bioinformatics analysis workflow.** An overview of the quality control steps and analyses done.

**Supplementary Fig. S2. Quality control metrics and expression of cell lineage markers across the spatial LUAD scRNA-seq dataset. A,** Statistical summary of cells passing quality control (QC) and showing cell number (left), fraction of mitochondrial genes (middle), and the number of detected genes (right) per sample. Dis, distant normal; Int, intermediate normal; Adj, adjacent normal; LUAD, tumor tissue. **B-C,** UMAP plots showing cells colored by patient ID (B) and library/sequencing batch (C). **D, F,** Bubble plots showing the percentage of cells expressing lineage markers (indicated by the size of the circle) as well as their scaled expression levels (indicated by the color of the circle) across all cells (D; related to main Fig. 1D and Fig. 1E) or selected cell types (F; related to main Fig. 1G and Fig. 1H). NK; natural killer cell, DC; dendritic cell, EC; endothelial cell. **E**, Stacked bar plots showing the relative cell fraction of spatial samples for major lineages considering non-proliferating cells (left) and proliferating ones (middle). Box plot showing fraction of proliferating epithelial cells in LUAD tissues versus normal spatial samples (right). *P -* value was calculated using Wilcoxon rank sum test.

**Supplementary Fig. S3. Analysis of clustering robustness of major cellular lineages**. **A,** UMAP plots showing cells colored by major lineages and identified with Harmony (left) or rPCA (middle). The right part of panel A shows a heatmap depicting the extent of cluster assignment overlap between rPCA (rows) and Harmony results (columns) as quantified by Jaccard index. **B**, UMAP plots (top) and the corresponding cluster overlap indices (bottom) when using 25% (left), 50% (middle), and 75% (right) of randomly sampled cells. The heatmaps show cluster overlap indices in randomly sampled results (rows) versus when using all 186,916 cells (columns).

**Supplementary Fig. S4. Transcriptome complexity of cells from P1. A-B,** Boxplots showing the number of detected transcripts (**A**) and the number of detected genes (**B**) in epithelial cells versus in all non-epithelial lineages including lymphoid, myeloid, and stromal cells. Boxplots showing the number of detected transcripts (**C**) and the number of detected genes (**D**) in proliferating epithelial cells versus in proliferating cells of non-epithelial lineages including lymphoid, myeloid, and stromal cells. Boxplots showing the number of detected transcripts (**E**) and genes (**F**) in non-proliferating epithelial cells versus in non-proliferating and non-epithelial lineages including lymphoid, myeloid, and stromal cells.

**Supplementary Fig. S5. Analysis of robustness of epithelial cell clustering. A,** UMAP view showing EPCAM+ cells colored by library/sequencing batch. **B,** UMAP plot showing EPCAM+ cells colored by patient ID. **C,** UMAP plots (top) showing clustering results when using 25% (top left), 50% (top middle), and 75% (top right) of randomly sampled epithelial cells. The corresponding heatmaps (bottom) show cluster overlap indices in randomly sampled results (rows) versus when using all 70,030 epithelial cells (columns).

**Supplementary Fig. S6. Cellular distribution of epithelial lineage clusters. A,** Bar plot showing absolute numbers of cells for each lung epithelial cell lineage. Dis, distant normal; Int, intermediate normal; Adj, adjacent normal; LUAD, tumor tissue; AT1, alveolar type 1; AT2, alveolar type 2. **B**, Stacked bar plot showing relative fractions of individual epithelial subclusters derived from each patient. **C.** Pie chart showing the fractional distribution of epithelial cells from the LUADs by epithelial lineage cluster. **D.** Box plot showing fraction of basal cells among epithelial cells and from LUADs versus other normal spatial samples. *P -* value was calculated using Wilcoxon rank sum test.

**Supplementary Fig. S7. Trajectory analysis of alveolar cells. A,** Potential developmental trajectory for alveolar cells inferred by pseudotime analysis. Cells were ordered by pseudotime (dotted box) and colored by alveolar cell state. **B,** Bubble plots showing the percentage (indicated by the size of the circle) of cells expressing markers of alveolar states shown in trajectory analysis from panel C as well as their scaled expression levels (indicated by the color of the circle) **C,** Pseudotime trajectory showing cells colored by Notch signaling signature score. **D,** Violin plots showing Notch signaling signature score among alveolar cell states in trajectory analysis from panel B.

**Supplementary Fig. S8. Inference of copy number variation (CNV) and by-patient subclustering of cells from the malignant-enriched cluster.** **A,** Heatmap showing CNV scores of cells in the malignant-enriched cluster. **B,** UMAP highlighting P2 cells among all patient cells in the malignant-enriched cluster (top left). Subclustering of P2 cells from malignant-enriched cluster showing (with red asterisk) cells harboring *KRAS*-G12D mutation (bottom left). Heatmap of top differentially expressed genes (DEGs) between cells of P2 malignant-enriched subclusters (right). UMAPs highlighting P3 (**C**), P4 (**D**) and P5 (**E**) cells among all patient subsets in the malignant-enriched cluster (top). UMAPs of by-patient subclustering of the malignant-enriched cluster (middle). Heatmaps of top DEGs between cells of by-patient malignant-enriched subclusters (bottom).

**Supplementary Fig. S9. Robustness of CNV inference in C9 cells from P3 and P5. A,** Heatmaps showing CNV scores of cells in the malignant-enriched cluster of P3 using NK cells (left), B cells (middle), T cells (right) as background references. **B,** Heatmaps showing CNV scores of cells in the malignant-enriched cluster of P5 using NK cells (left), B cells (middle), T cells (right) as background references.

**Supplementary Fig. S10. Phylogenetic reconstruction analysis using inferred large-scale CNVs in P3.** Hierarchical clustering tree of P3 copy number profile (top) and the phylogenetic tree of the same profile (bottom). *P* – value was calculated using the Pearson’s correlation test.

**Supplementary Fig. S11. Quality control metrics of cells from lymphoid cell subsets. A-B,** UMAP of lymphoid cells colored by patient ID (A) and sequencing/library batch (B). **C,** Bar plots showing the absolute numbers of each lymphoid cell subset in each spatial sample. Dis, distant normal; Int, intermediate normal; Adj, adjacent normal; LUAD, tumor tissue; CTL, cytotoxic T lymphocyte; Treg, T regulatory cell; ILC, innate lymphoid cell; NK, natural killer cell.

**Supplementary Fig. S12. Analysis of robustness of CD8+ T cell clustering.** UMAP plots (top) showing the clustering results when using 25% (top left), 50% (top middle), and 75% (top right) of randomly sampled CD8+ T cells. The corresponding heatmaps (bottom) show cluster overlap indices in randomly sampled results (rows) versus the original result using all 9,274 CD8+ T cells (columns).

**Supplementary Fig. S13. Reprogramming of lymphoid cell subsets towards an immune suppressive tumor microenvironment in early-stage LUAD. A,** Changes in the abundance of specific lymphoid cellular lineages and states across the LUADs and spatial normal samples. Embedded pie charts show the contribution of each spatial sample to the indicated cell subtype/state. **B,** Cytotoxicity signature score in CD8+ *GNLY*_hi CTLs in P3 (left) and in P4 (right) across spatial samples (*, *P* < 0.05; ***, *P* < 0.001 of the Wilcoxon rank sum test). **C,** Treg signature score in T regulatory cells across spatial samples in all patients (left), P3 (middle) and in P5 (right) (*, *P* < 0.05; **, *P* < 0.01; ***, *P* < 0.001 of the Wilcoxon rank sum test). **D,** Depletion of CD4+ CTLs in the tumor microenvironment of LUAD. Boxplot showing percentage of CD4+ CTLs *GZMA*-hi among total CD4+ cells from all patients across the spatial samples. Each circle represents a patient sample. *P -* value was calculated using Kruskal-Wallis test. **E,** Violin plots showing cytotoxic signature score in CD4+ CTL *GZMA*-hi cells from all patients (left) and P5 (right; ***, *P* < 0.001 of the Wilcoxon rank sum test). **F,** Frequency of CD4+ CTL *GZMA***-**hi cells co-expressing *GZMA* and *GZMH* across the spatial samples. The fractions of *GZMA+GZMH+ CD4+* CTLs are labeled on each plot*.* **G-I,** Expression profiling of different isotypes of plasma cells across the spatial samples. **G,** Heatmap showing isotype genes expression in plasma cells. **H,** Bar plots showing plasma cell isotype composition across spatial samples. **I,** Dot plots showing the fractional change of IGHA1/2 and IGHG3 across all patients and in each patient (left to right). **J-M,** Reprogramming of B cells in early-stage LUAD. **J,** Heatmap showing unsupervised clustering of B cells sub-populations. **K,** UMAP showing re-clustering of B cells into sub-populations. **L,** Ridge plots (left) showing *RAC2* and *ACTG* expression levels among B cell sub-clusters, and scatter plots (right) showing the frequency of B cells co-expressing *RAC2+* and *ACTG*+ across spatial samples. The fractions of *RAC2+ACTG*+ B cells are labeled on each plot*.* **M,** Boxplot showing the B cell C0 signature score in normal lung tissues (NL), premalignant atypical adenomatous hyperplasias (AAH) and LUADs in an independent validation cohort. (*, *P* < 0.05; ***, *P* < 0.001; N.S, *P* > 0.05 of the Wilcoxon rank sum test).

**Supplementary Fig. S14. Changes in fractions of lymphoid cell subsets in association with smoking status. A,** Stacked bar plot showing changes in the relative fractions of lymphoid cell subsets between non-smokers and smokers. Treg, T regulatory cell; ILC, innate lymphoid cell; NK, natural killer cell; Other T cells, other CD4 and CD8 T cells states. **B**, Boxplot showing change in relative fraction of plasma cells between non-smoker and smokers in TCGA LUAD cohort. *P -* value was calculated using Wilcoxon rank sum test.

**Supplementary Fig. S15. Association of a B cell signature with clinical outcome.** Overall survival (OS, left) and progression-free interval (PFI, right) of treatment-naïve LUAD patients grouped by scores of the B cell signature in TCGA (**A**) and MDACC cohorts (**B**). Patients were stratified based on the first and fourth quartiles of B cell signature scores (TCGA B cell low n = 116 and high n = 114; MDACC B cell low n = 14 and high n = 14). Survival analysis was performed using Kaplan–Meier estimates and two-sided log-rank tests.

**Supplementary Fig. S16. Quality control metrics of cells from myeloid cell subsets. A-B,** UMAP view showing myeloid cells colored by patient ID (A) and library/sequencing batch (B). **C**, Stacked bar plots showing absolute numbers of each myeloid cell subset across the spatial samples. Dis, distant normal; Int, intermediate normal; Adj, adjacent normal; LUAD, tumor tissue; mono; monocytes, mac; macrophages, cDC; classical dendritic cells, pDC; plasmacytoid dendritic cell.

**Supplementary Fig. S17. Reprogramming of myeloid cells in the tumor microenvironment of early-stage LUAD. A,** Changes in the abundance of specific myeloid cellular lineages and states across the LUADs and spatial normal samples. Embedded pie charts show the contribution of each spatial sample to the indicated cell subtype/state. **B**, Boxplot showing the antigen presentation signature score between M2-like Mac_c1 and M2-like Mac_c5. Mac; macrophages. *P -* value was calculated using Wilcoxon rank sum test. **C**, Violin plots showing the expression level changes among spatial samples for antigen presentation involving genes. **D**, Heatmap showing the expression of inflammatory and non-inflammatory signature genes in cDC2 cells. **E**, Boxplot showing the non-inflammatory versus inflammatory signature score in normal lung tissues (NL), premalignant atypical adenomatous hyperplasias (AAH) and LUADs in an independent validation cohort (**, *P* < 0.01; ***, *P* < 0.001; N.S, *P* > 0.05 of the Wilcoxon rank sum test). **F**, Heatmap showing DEGs between LUAD and normal samples in pDCs. **G**, Ridge plots showing the expression level changes of *FOS*, *JUN* and *FOSB* in pDCs and across spatial samples.

**Supplementary Fig. S18. Analysis of robustness of monocyte and macrophage clustering.** UMAP plots (top) showing the Harmony-based clustering results when using 25% (top, left), 50% (top, middle), and 75% (top, right) of randomly sampled cells. The corresponding heatmaps (bottom) show cluster overlap indices in randomly sampled results (rows) versus when using all 27,664 monocytes and macrophages (columns).

**Supplementary Fig. S19. Survival analysis based on dendritic cell signatures.** OS (left) and PFI (right) of treatment-naïve LUAD patients grouped by the cDC2 signature scores (ratio of non-inflammatory to inflammatory expression) in TCGA (**A**) and the MDACC (**B**) cohorts. Patients were grouped into first and fourth quartiles based on the cDC2 signature scores (TCGA cDC2 low n = 113 and high n = 116, MDACC cDC2 low n = 14 and high n = 14). Survival analysis was performed using Kaplan–Meier estimates and two-sided log-rank tests.

**Supplementary Fig. S20. Composition and gene expression changes in stromal and endothelial sub-populations in early-stage LUAD. A,** UMAP view of stromal and endothelial cells colored by sample batch. **B**, UMAP view of stromal and endothelial sub-populations. **C**, Bubble plot showing the percentage of stromal and endothelial cells expressing lineage markers (indicated by the size of the circle) as well as their scaled expression levels (indicated by the color of the circle). **D,** Bar plot showing the absolute number of cells from each stromal and endothelial subset. Dis, distant normal; Int, intermediate normal; Adj, adjacent normal; LUAD, tumor tissue; EC, endothelial cells. **E**, Changes in the abundance of stromal and endothelial cell lineages across the LUADs and spatial normal samples. Embedded pie charts show the contribution of each spatial sample to the indicated stromal and endothelial subtype. **F**, Heatmap showing DEGs between LUAD and normal samples in EC venule sub-populations. **G**, Bar plot showing significantly enriched pathways of up/down regulated DEGs in panel F. **H,** Ridge plots showing the expression level changes of *HLA-DPB1*, *IL33*, and *IGFBP7* in EC venule sub-populations and across spatial samples.

**Supplementary Fig. S21. Spatially enriched chemokine-receptor cell-cell communication networks in the immune microenvironment of early-stage LUAD. A,** Circos plot showing modulation of chemokine-mediated ligand-receptor (L-R) pairs in the LUAD relative to tumor-intermediate normal tissue of P3. **B,** Violin plots showing spatial expression levels of the major chemokine and receptor genes in P3. **C,** Circos plots showing modulation of chemokine-mediated L-R pairs between the LUADs and tumor-intermediate or -distant normal tissues of P2 (left) and P5 (right), respectively. **D**, Violin plots showing spatial expression levels of the major chemokine and receptor genes in P5.

**Supplementary Fig. S22. Single-cell expression patterns of *CD24* in the ecosystem of early-stage LUAD. A,** Violin plot showing *CD24* expression levels across major lineages from all patient samples**. B,** Violin plots showing *CD24* expression levels across major lineages for each patient.

**Supplementary Fig. S23. Correlation of *CD24* expression with immune genes.** Scatterplots showing gene expression correlation analysis between *CD24* and immune genes in a cohort of normal lung (NL) tissues, premalignant atypical adenomatous hyperplasias (AAH), and invasive LUADs (GSE10251, **A**) and in LUADs and NL from TCGA cohort (**B**). *P* - values were calculated using Pearson’s correlation test.

**Supplementary Fig. S24. CRISPR-mediated knockdown of *Cd24a* expression in mouse LUAD cells.** Expression of *Cd24a* in MDA-F471 mouse LUAD cells transduced with sgCt or sg*Cd24a* CRISPR constructs and following FACS-sorting for high or low expression of CD24 surface protein, respectively. Gene expression was normalized to both *Gapdh* and *Actinb* using the 2^-ΔΔCt^ method (***, *P* < 0.001; unpaired Student’s *t* test).
