## Supplementary Figures for "Resolving the spatial and cellular architecture of lung adenocarcinoma by multi-region single-cell sequencing"

### Supplementary Fig. S1

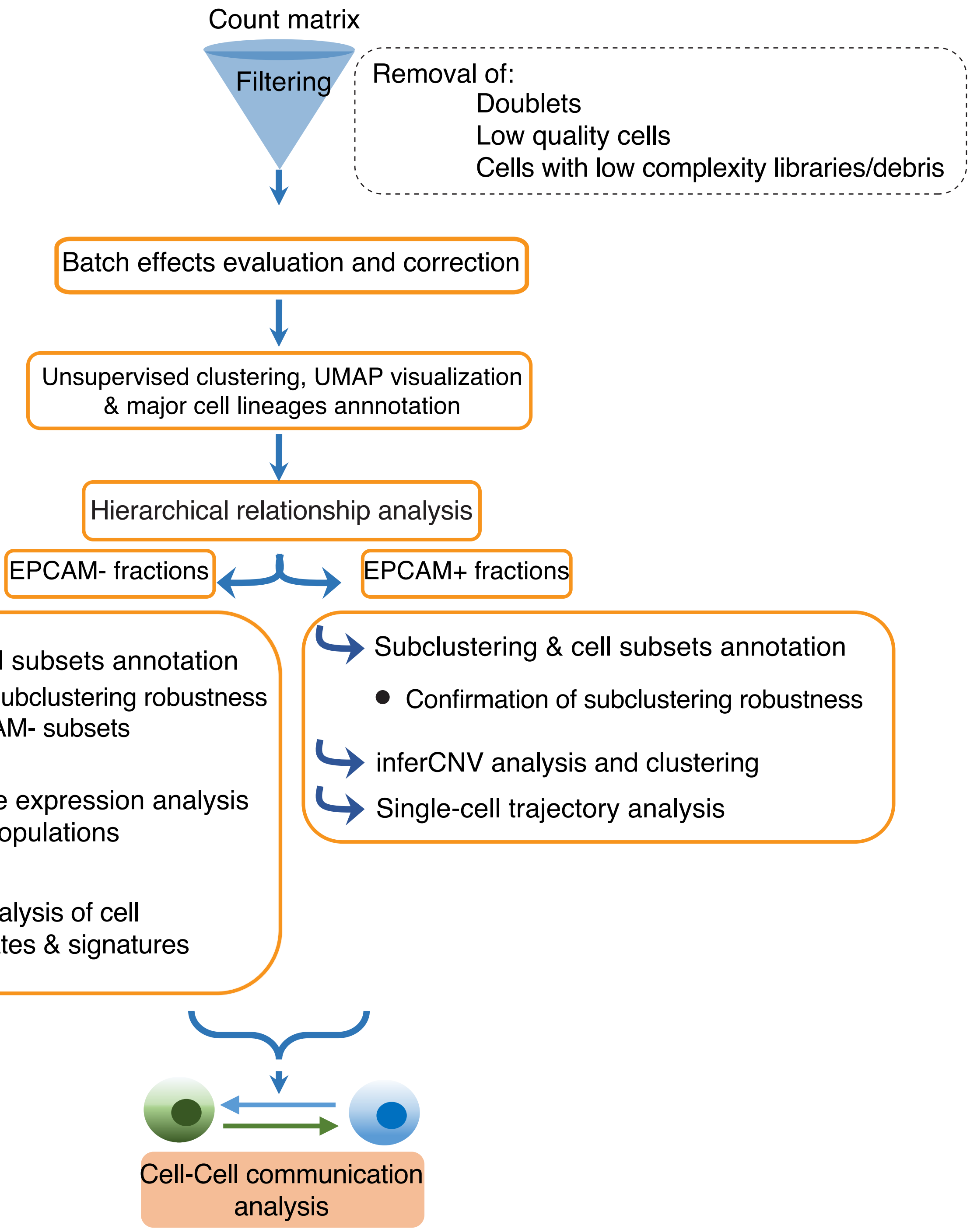

### Supplementary Fig. S2

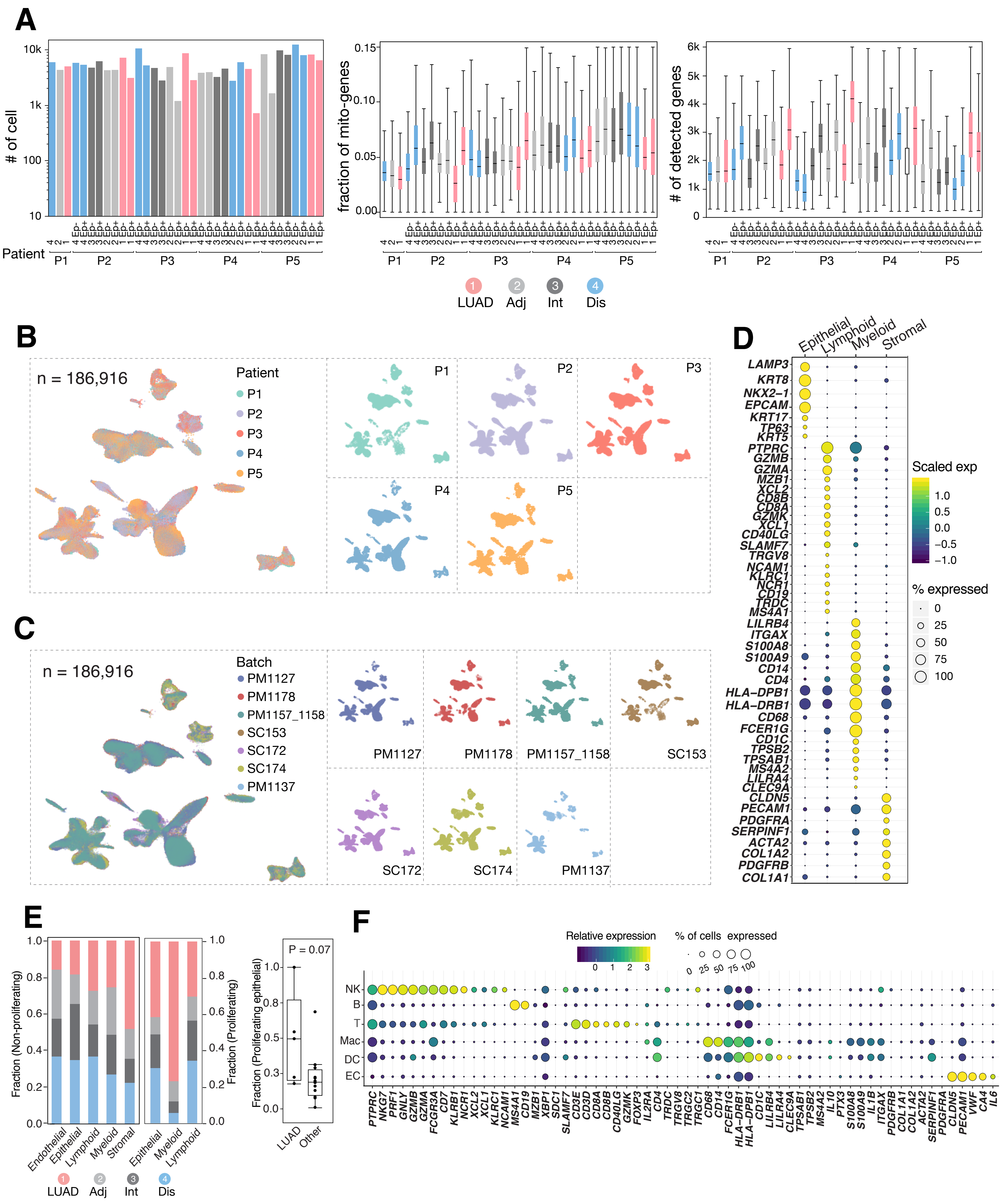

### Supplementary Fig. S3

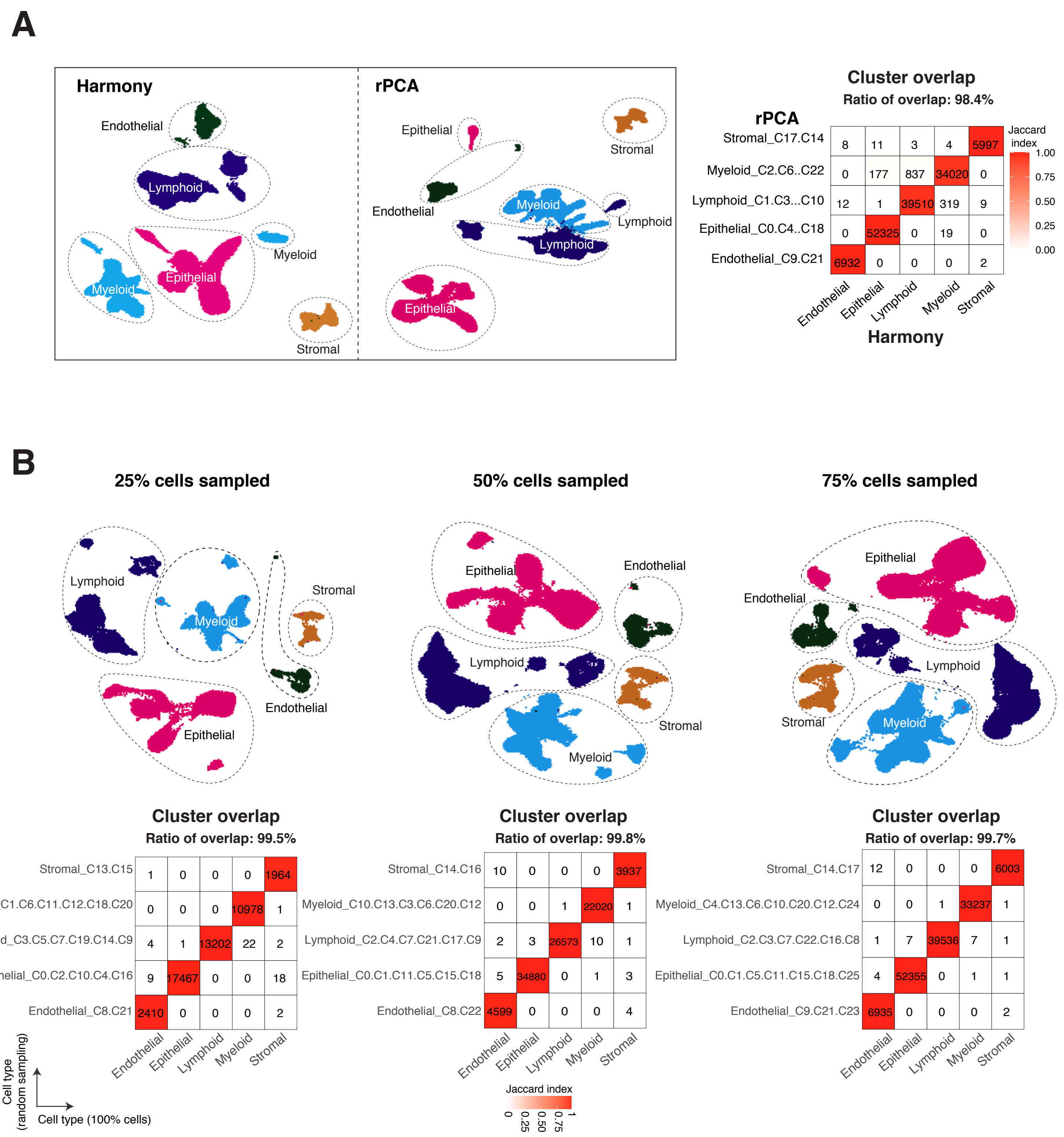

### Supplementary Fig. S4

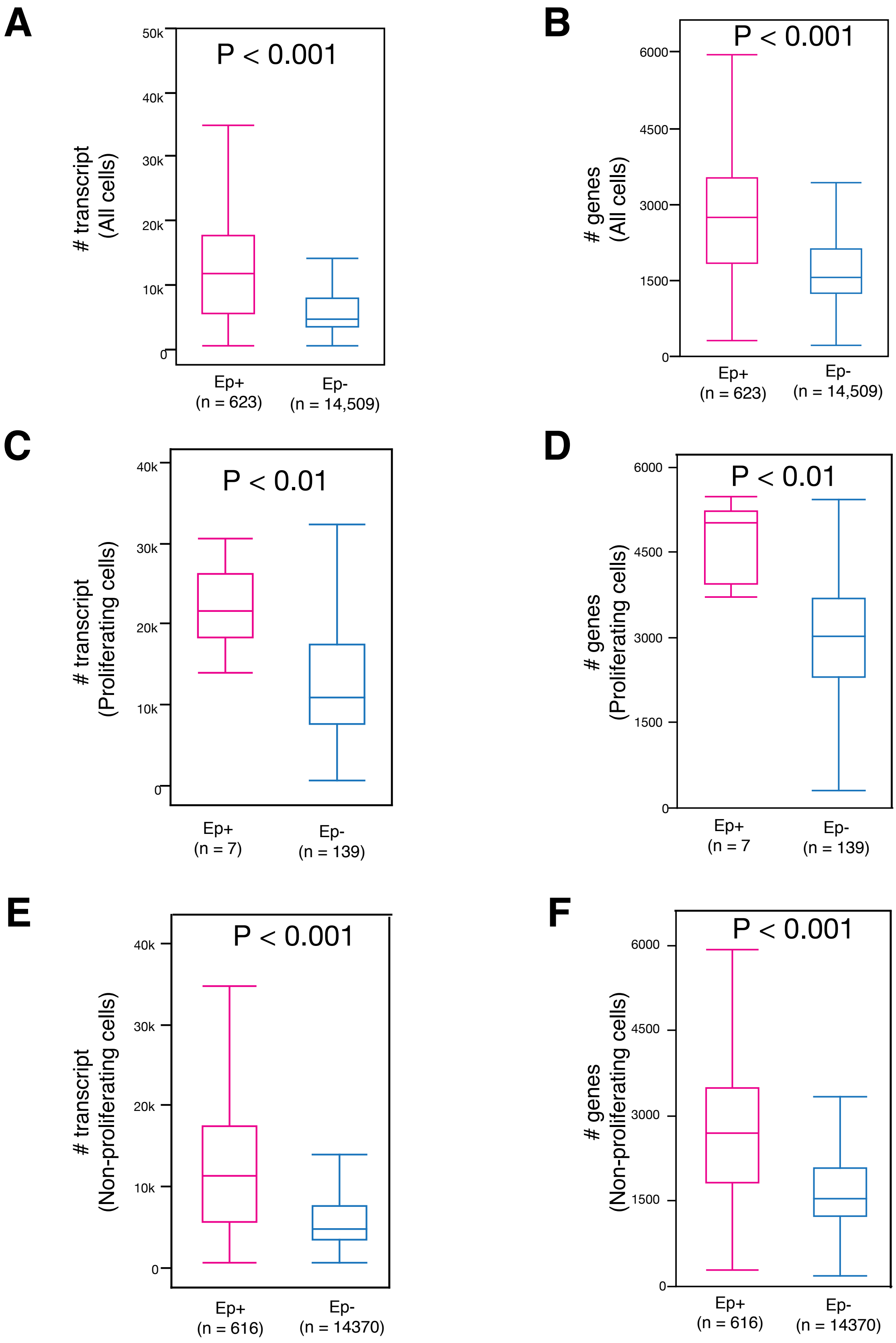

### Supplementary Fig. S5

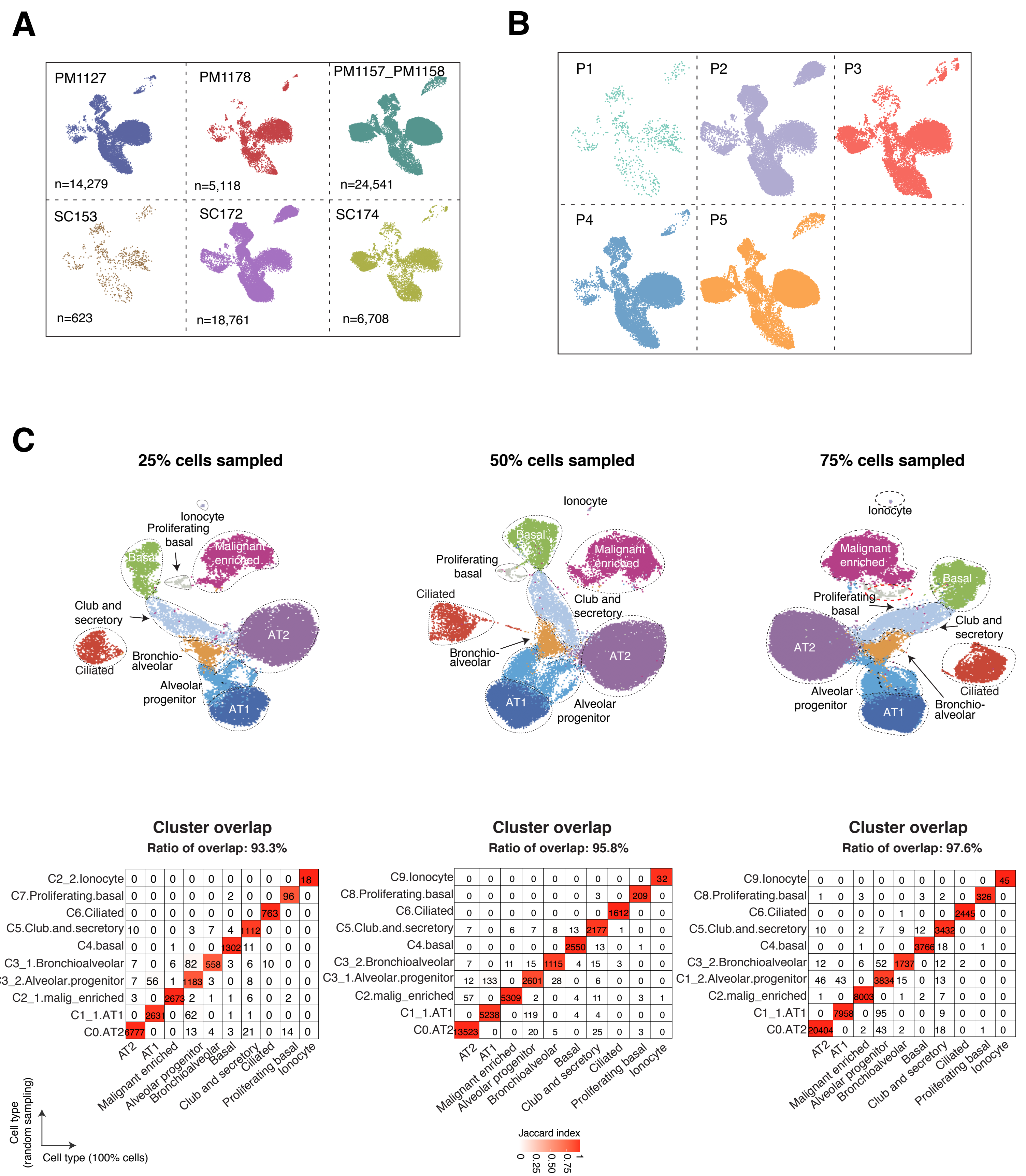

### Supplementary Fig. S6

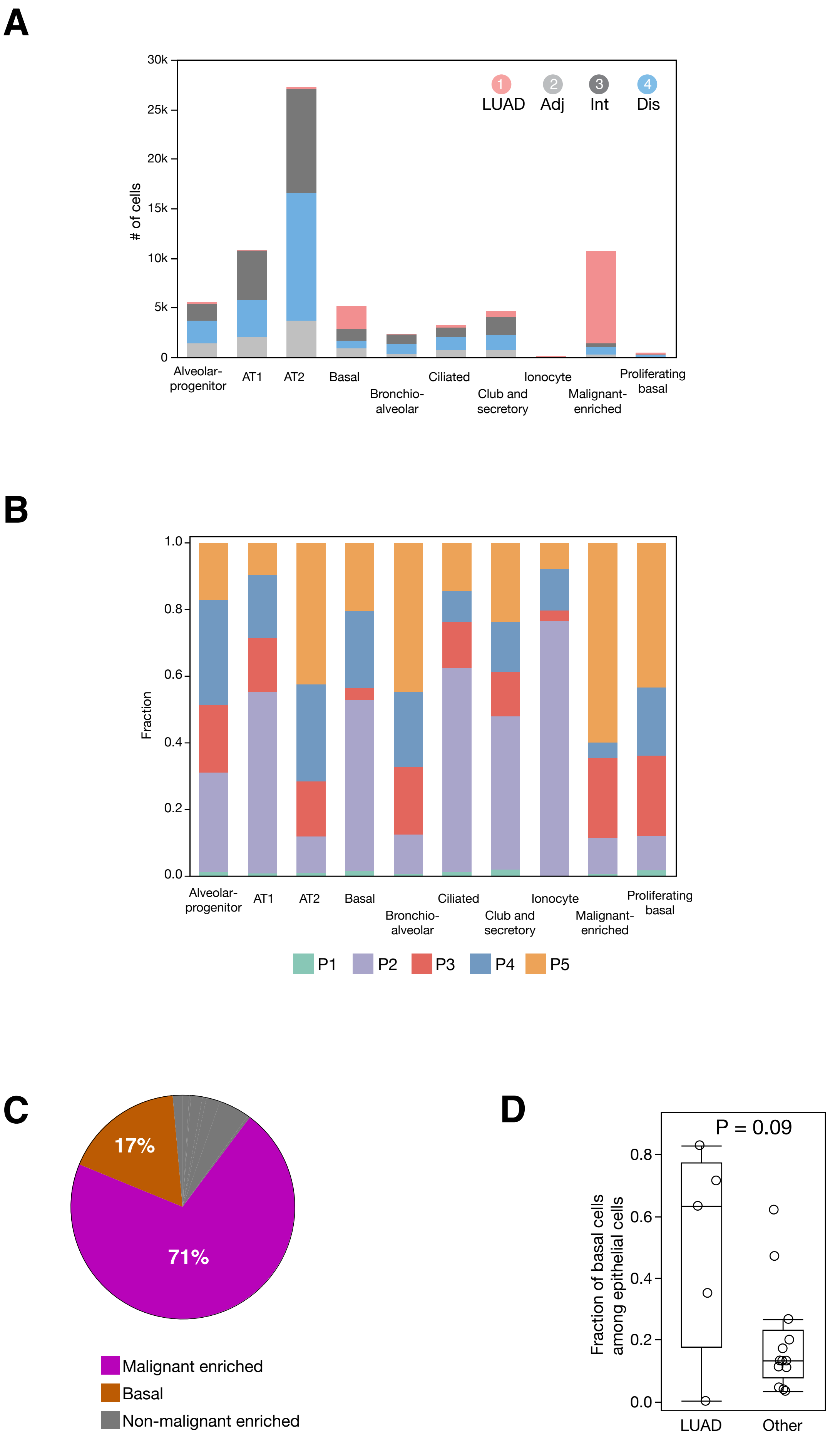

### Supplementary Fig. S7

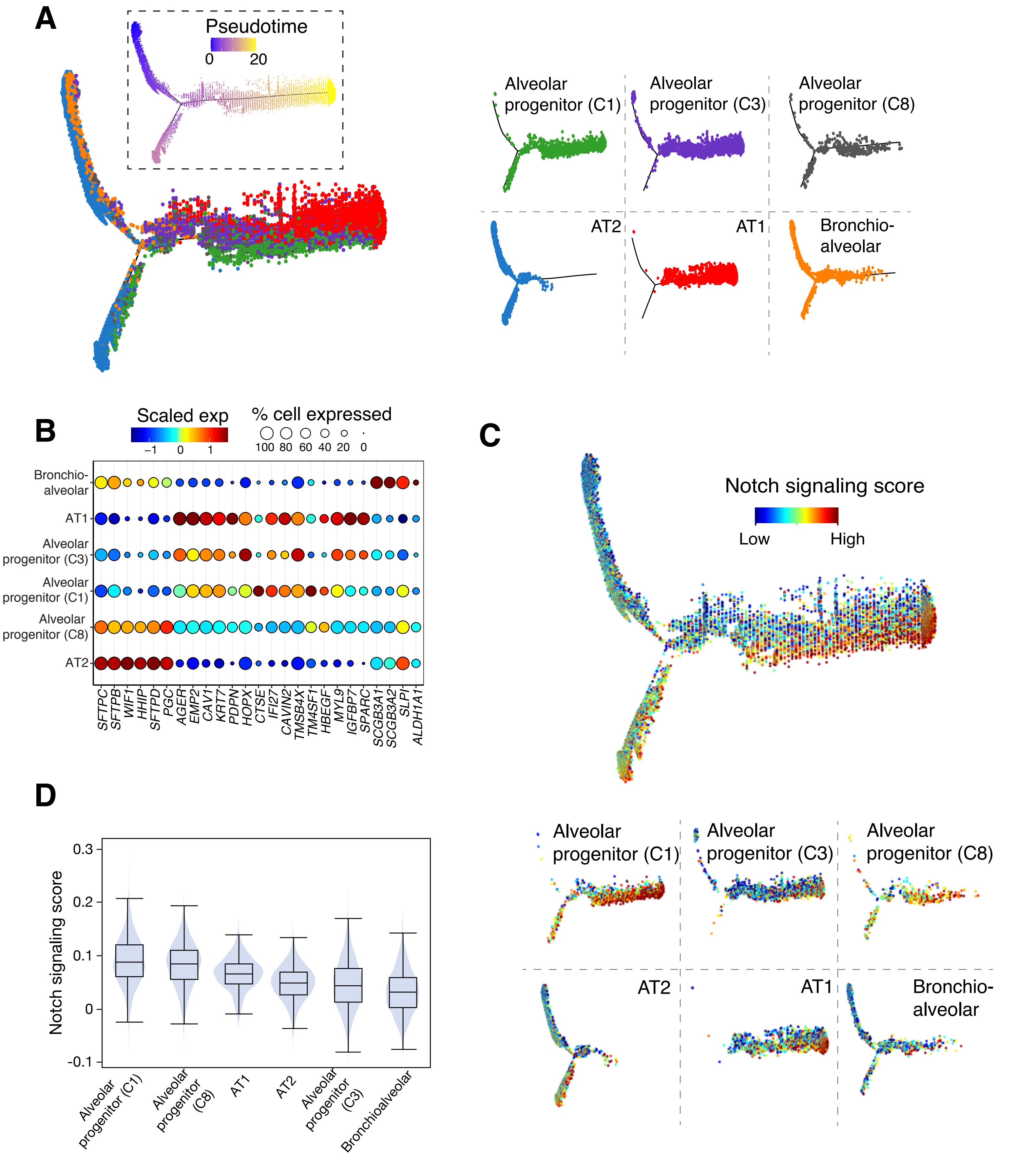

### Supplementary Fig. S8

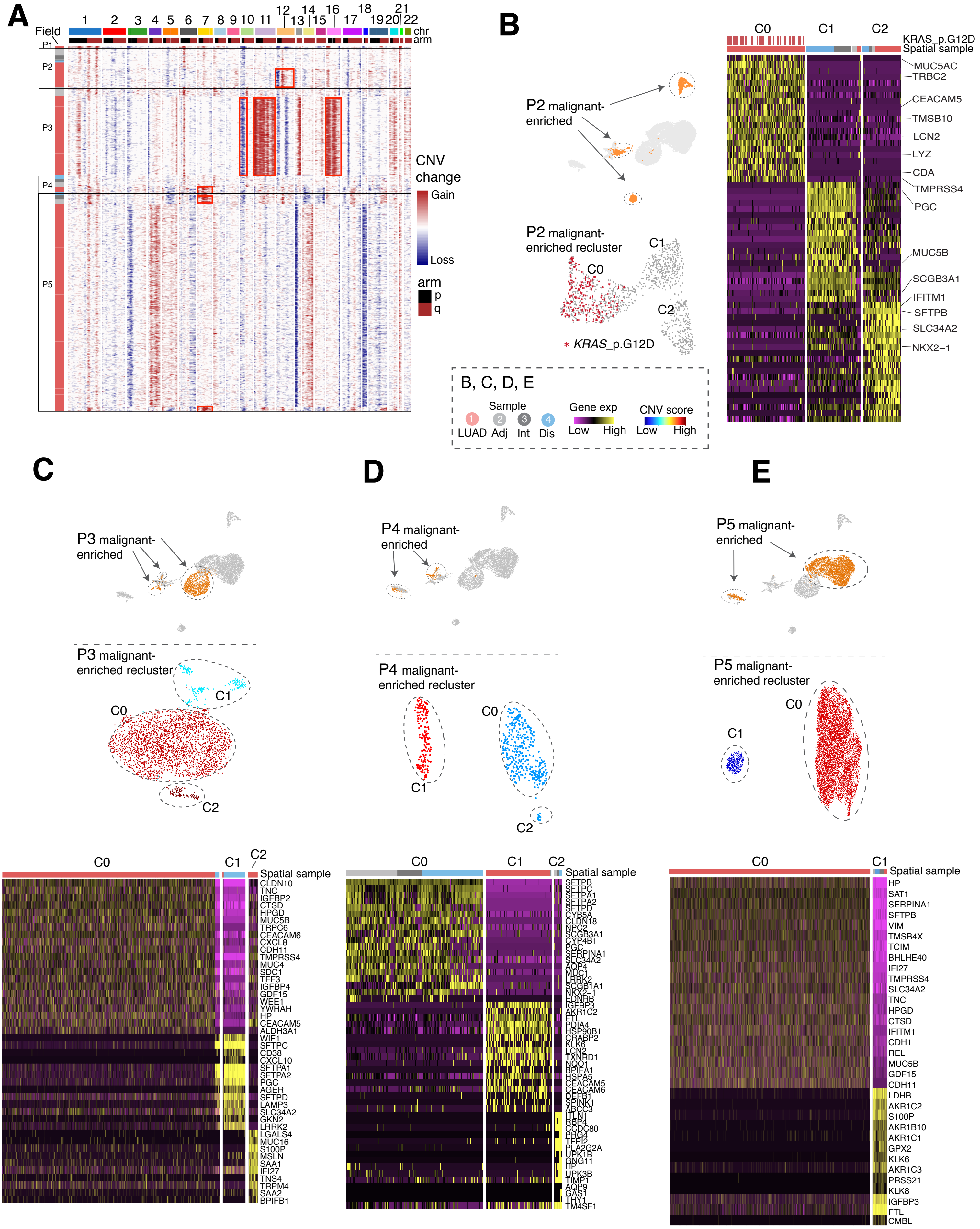

### Supplementary Fig. S9

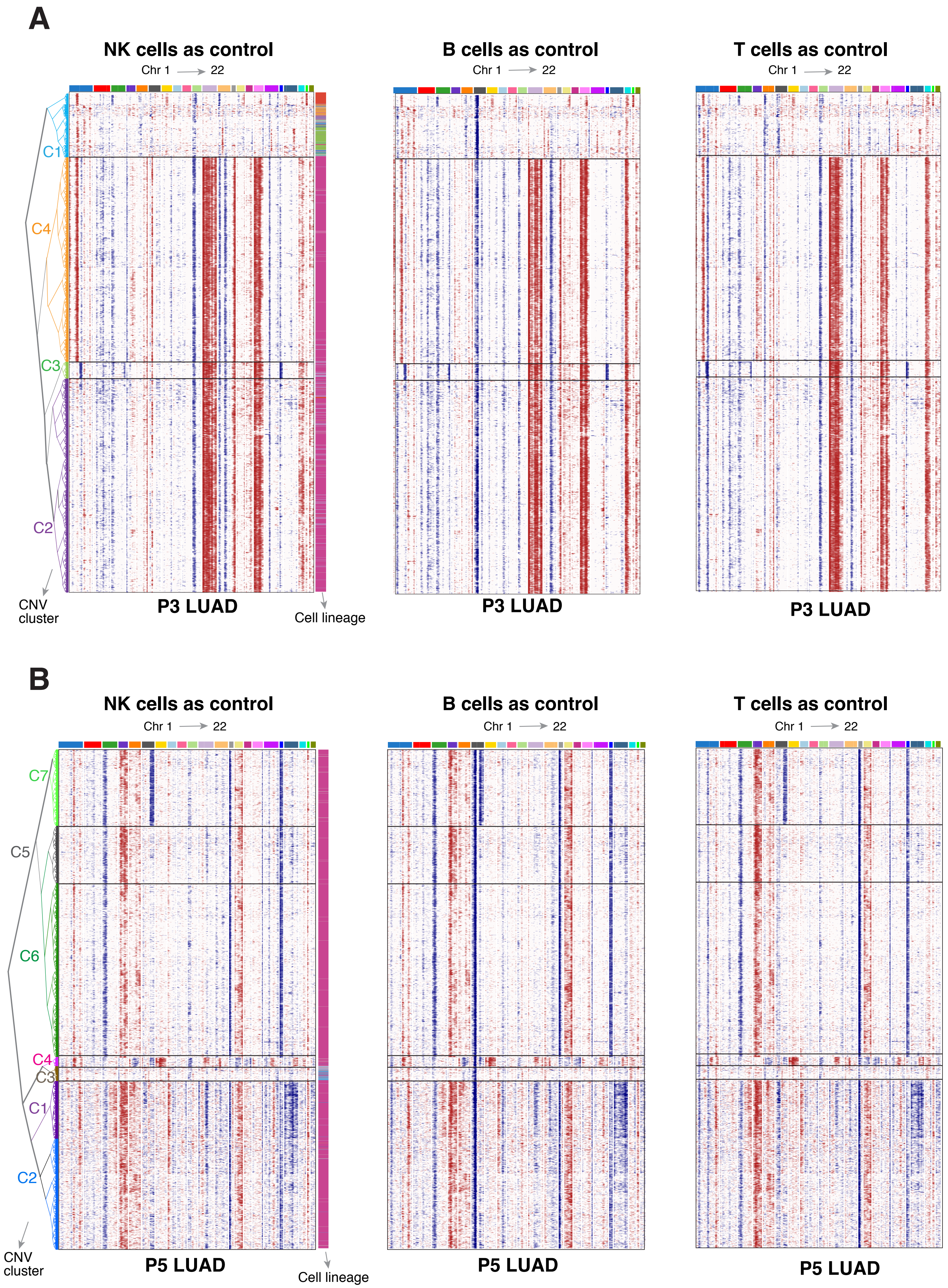

### Supplementary Fig. S10

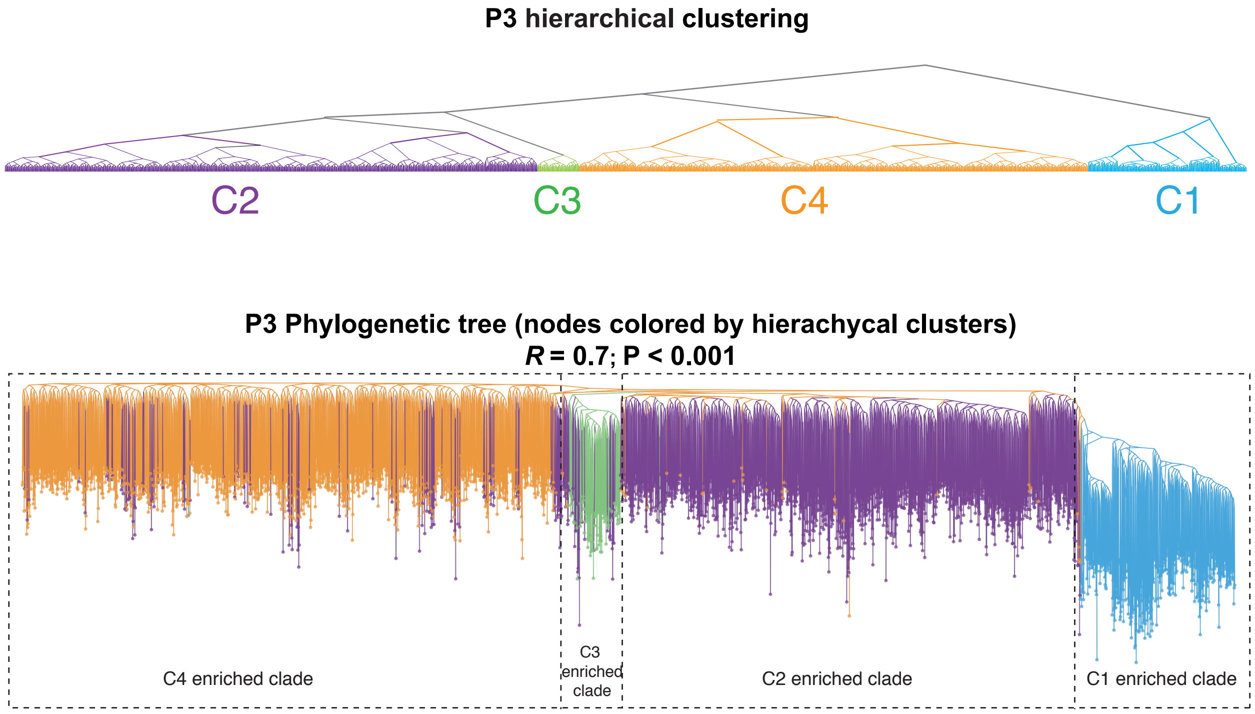

### Supplementary Fig. S11

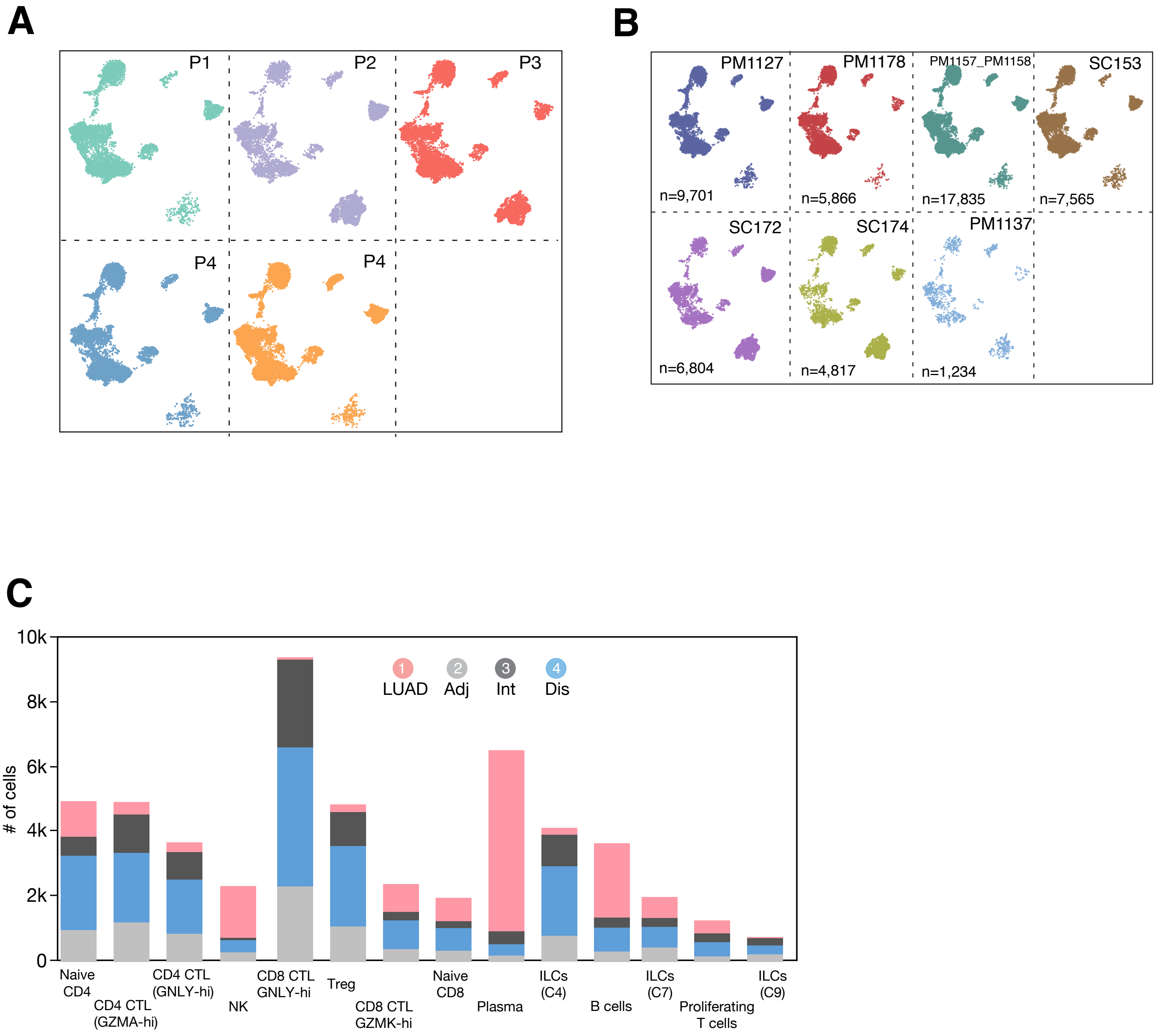

### Supplementary Fig. S12

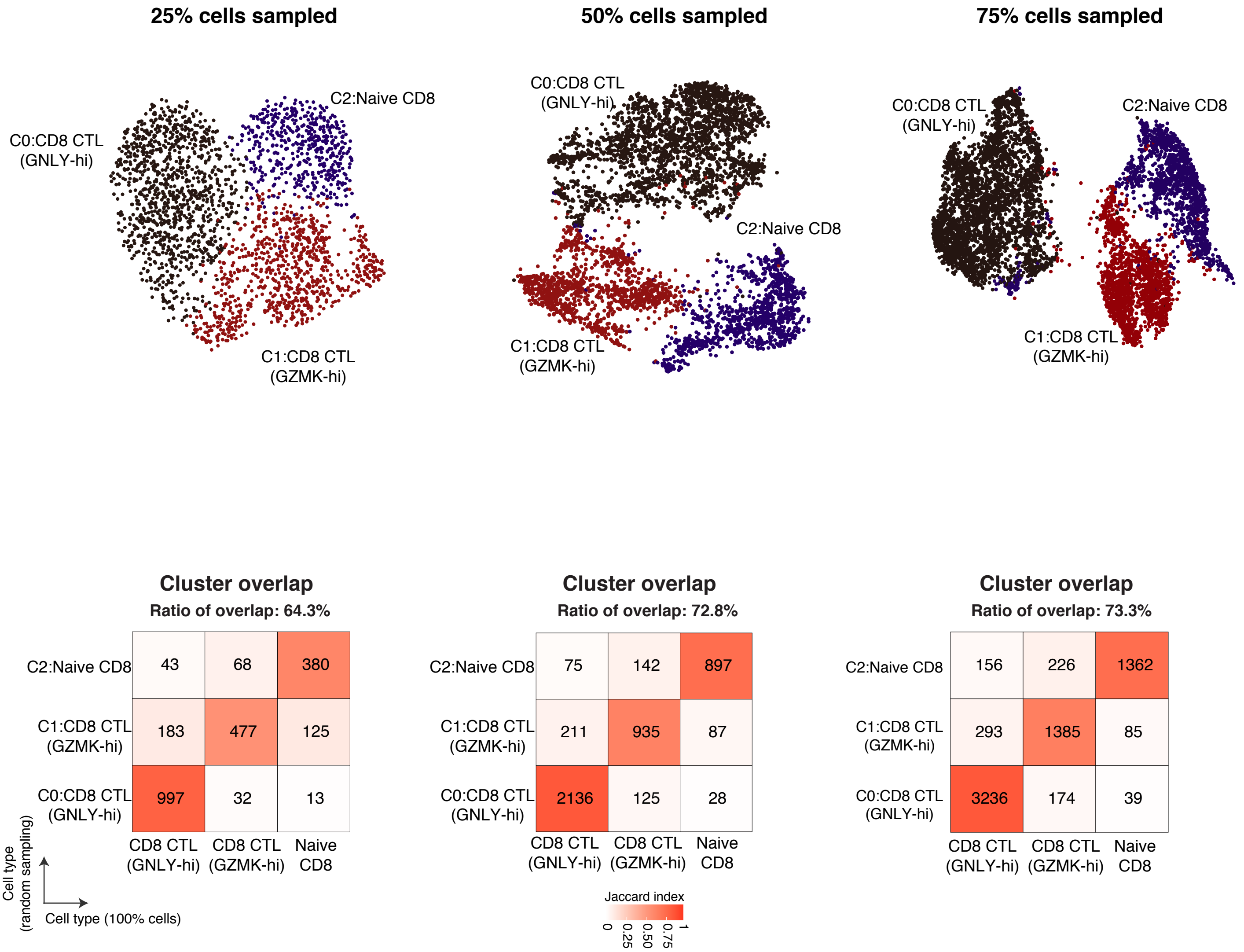

### Supplementary Fig. S13

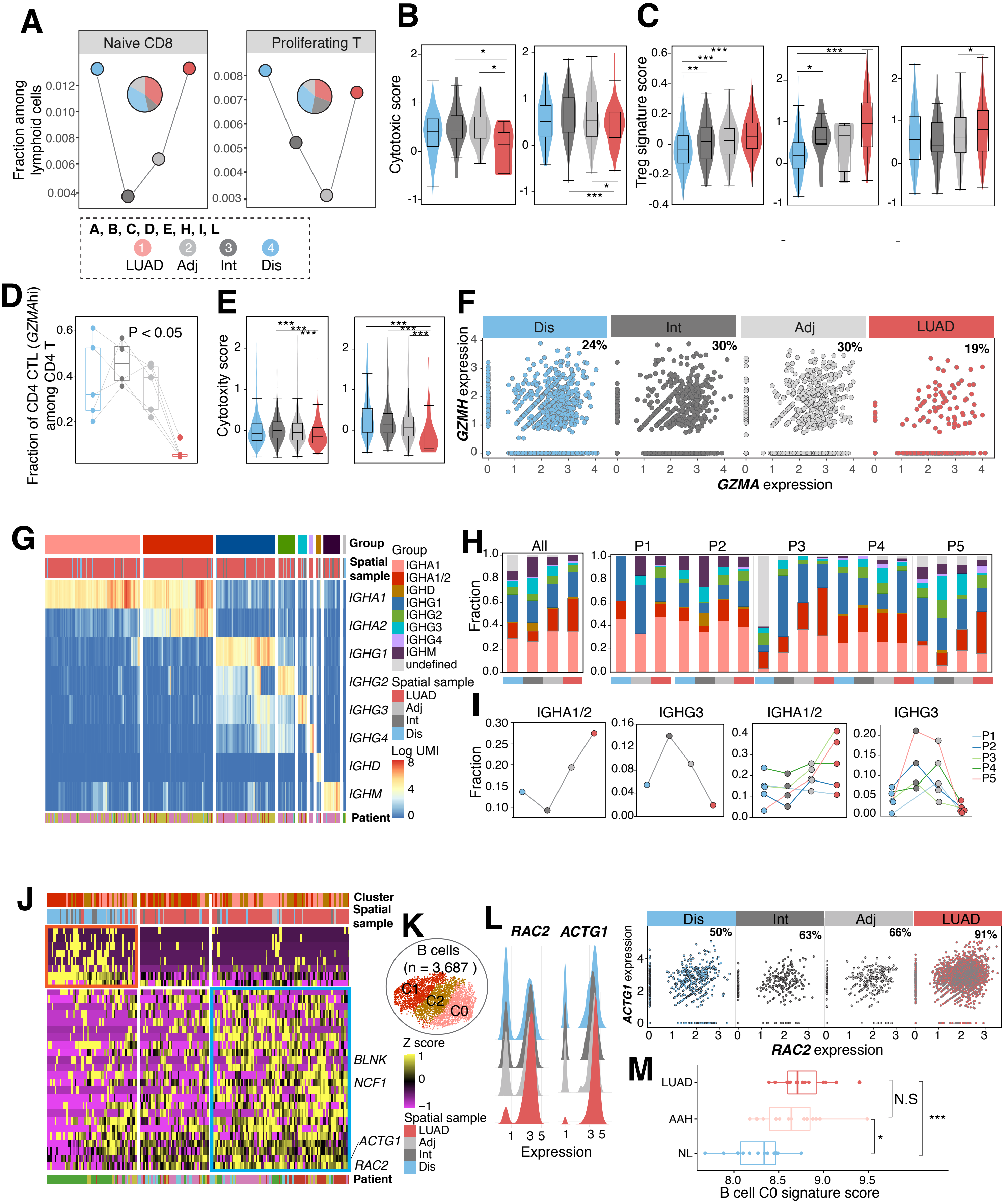

### Supplementary Fig. S14

A

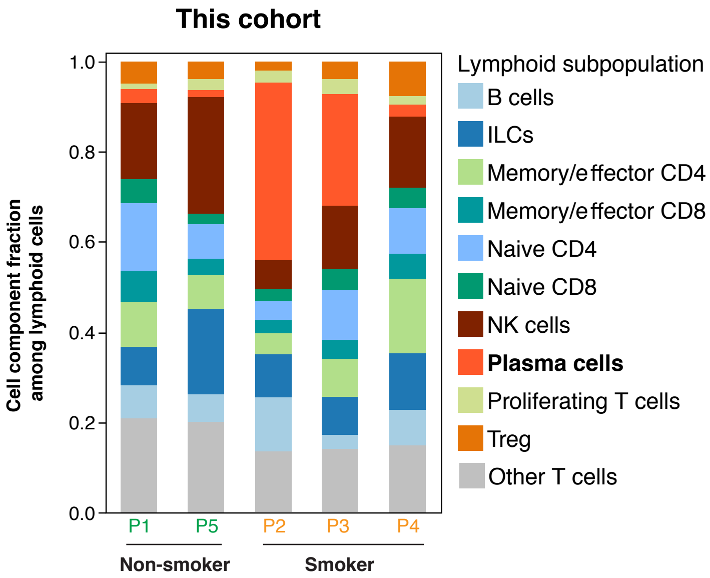


B

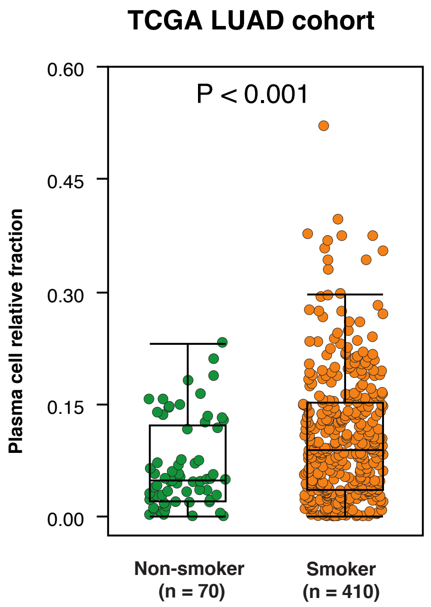

### Supplementary Fig. S15

**A**

B cell signature, TCGA LUAD cohort

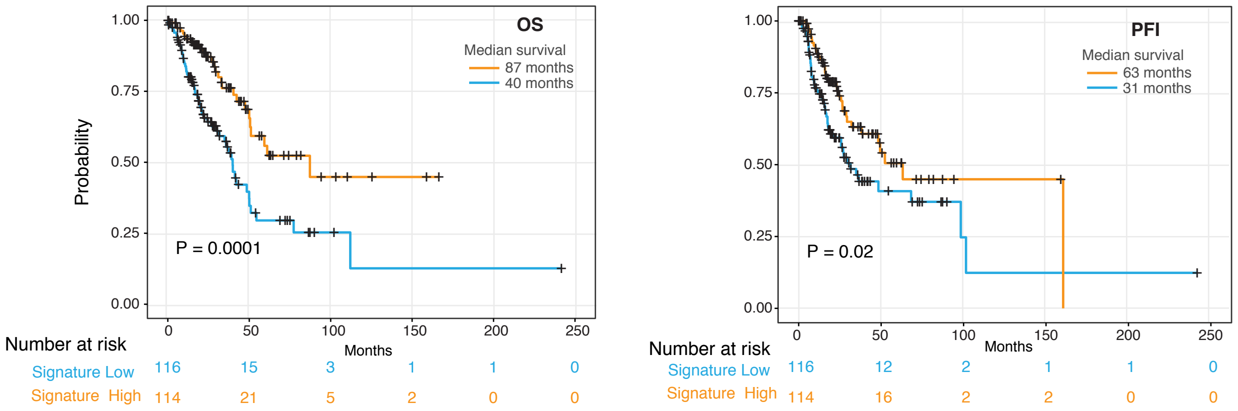


**B**

B cell signature, MDACC cohort

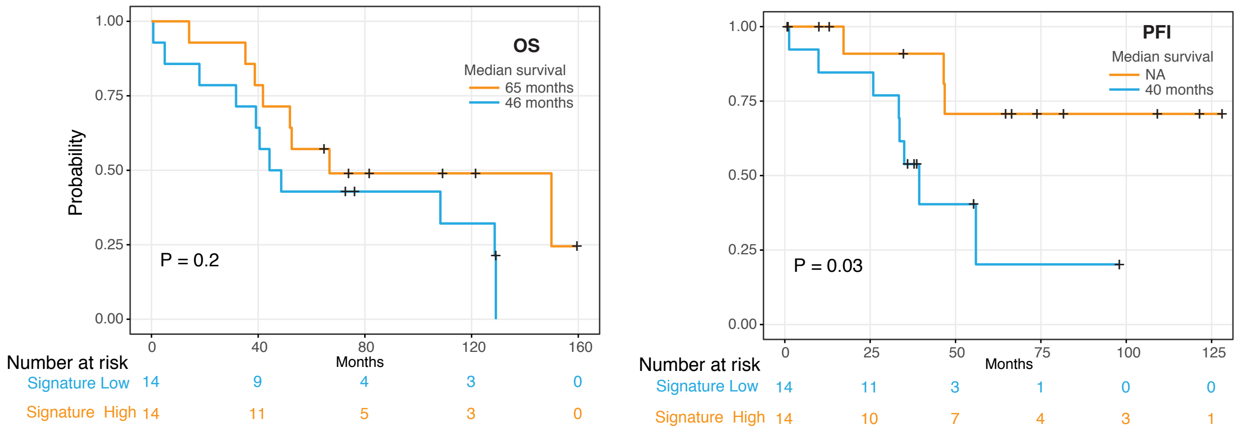

### Supplementary Fig. S16

A

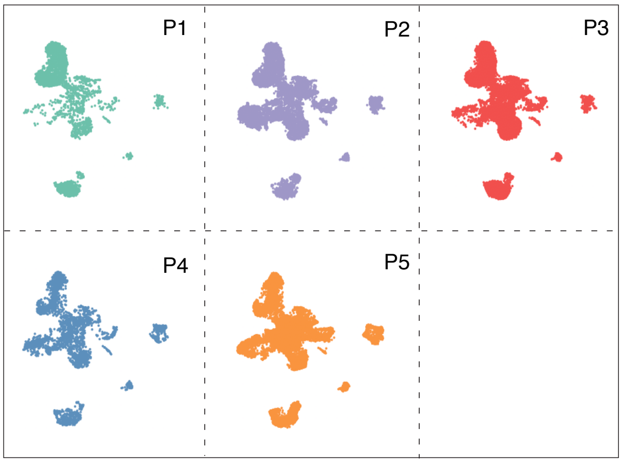


B

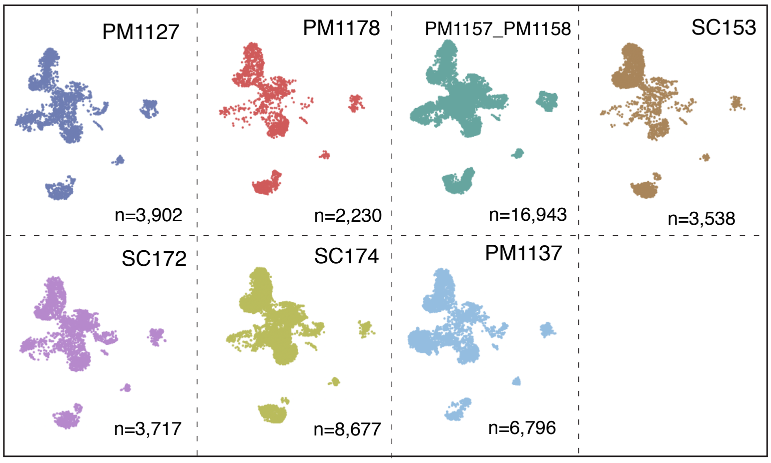


C

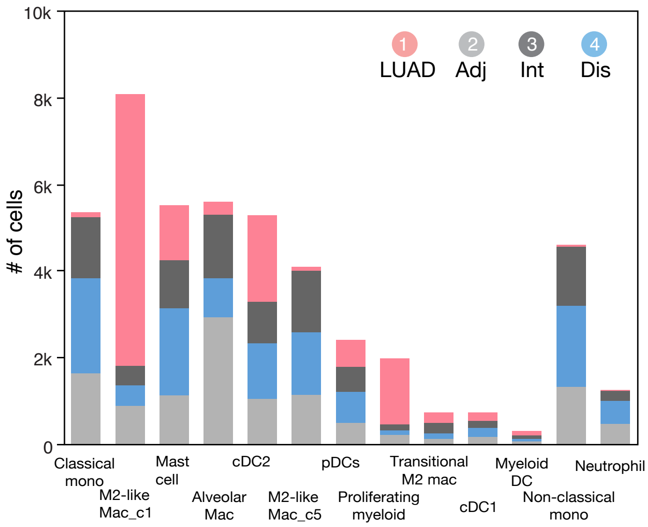

### Supplementary Fig. S17

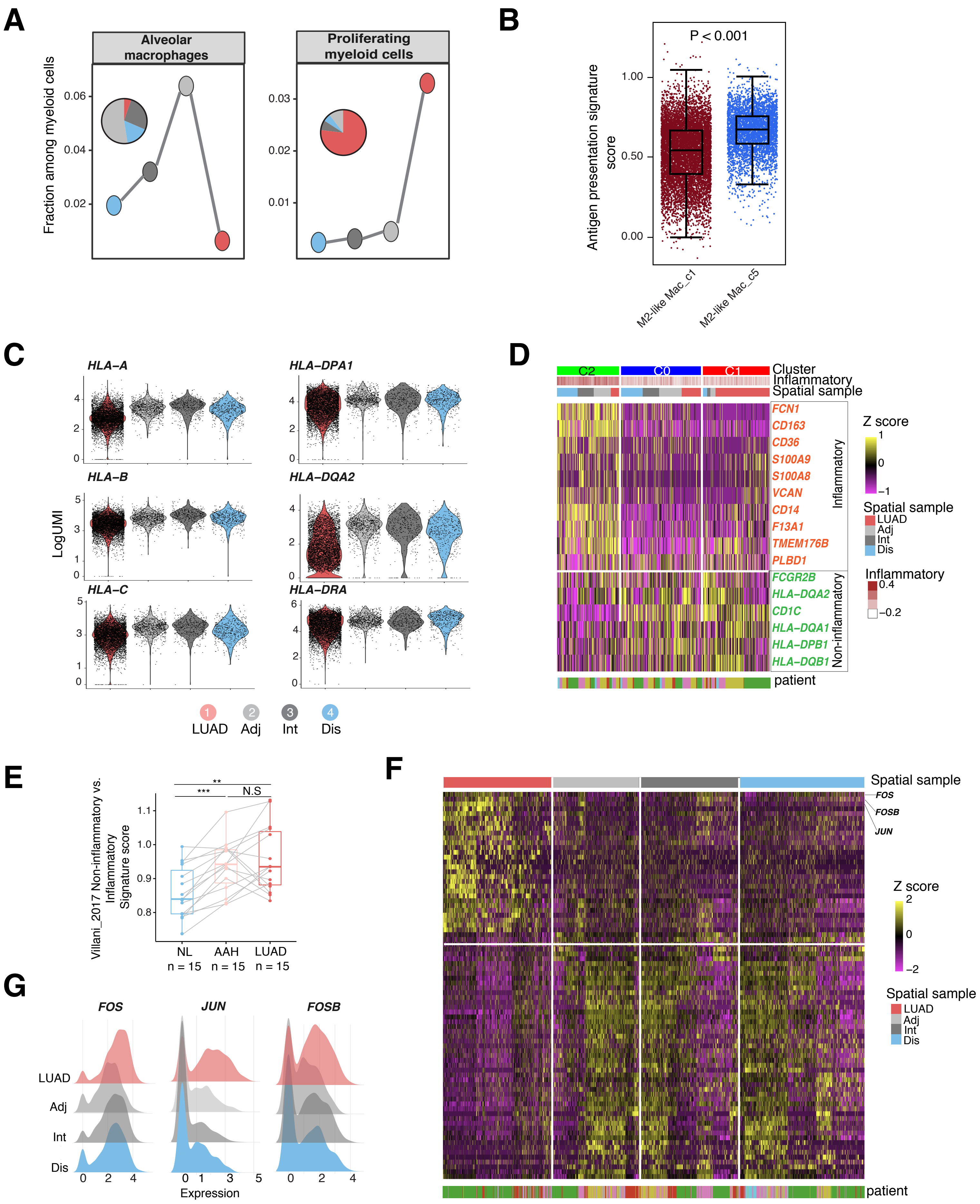

### Supplementary Fig. S18

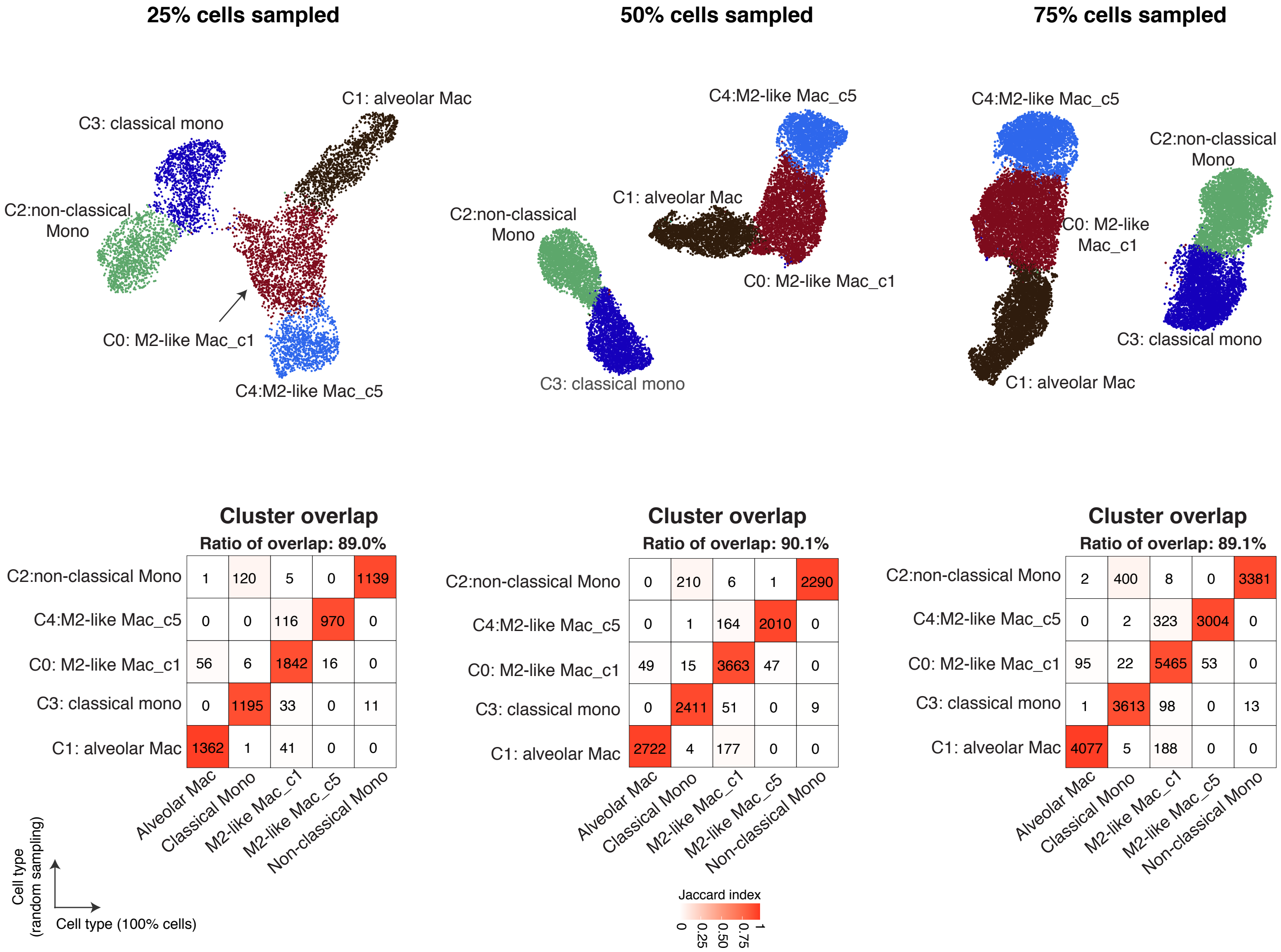

### Supplementary Fig. S19

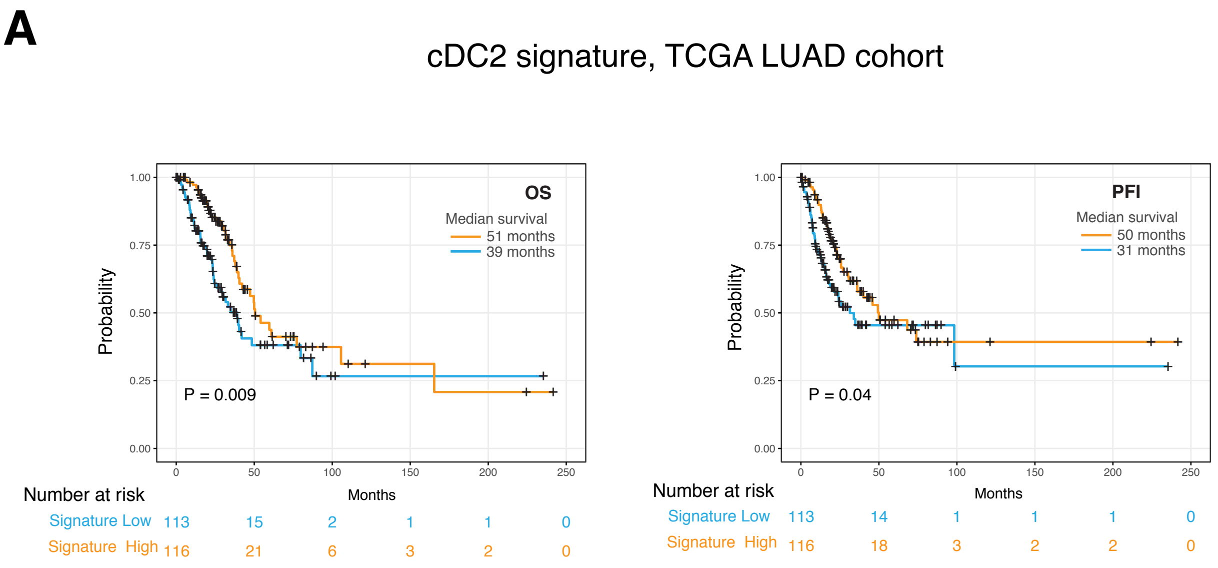


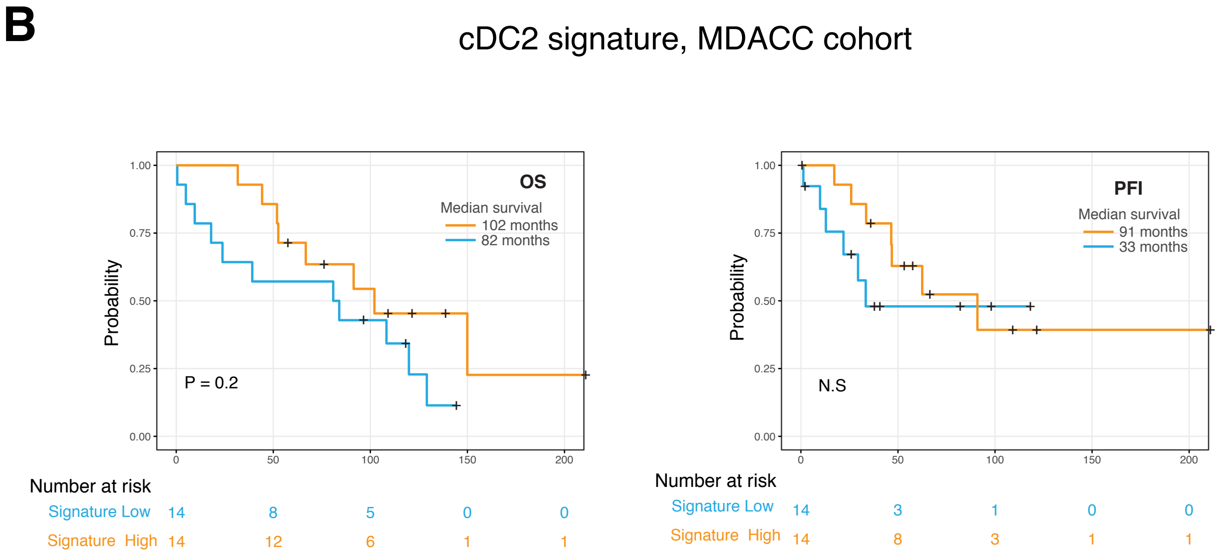

### Supplementary Fig. S20

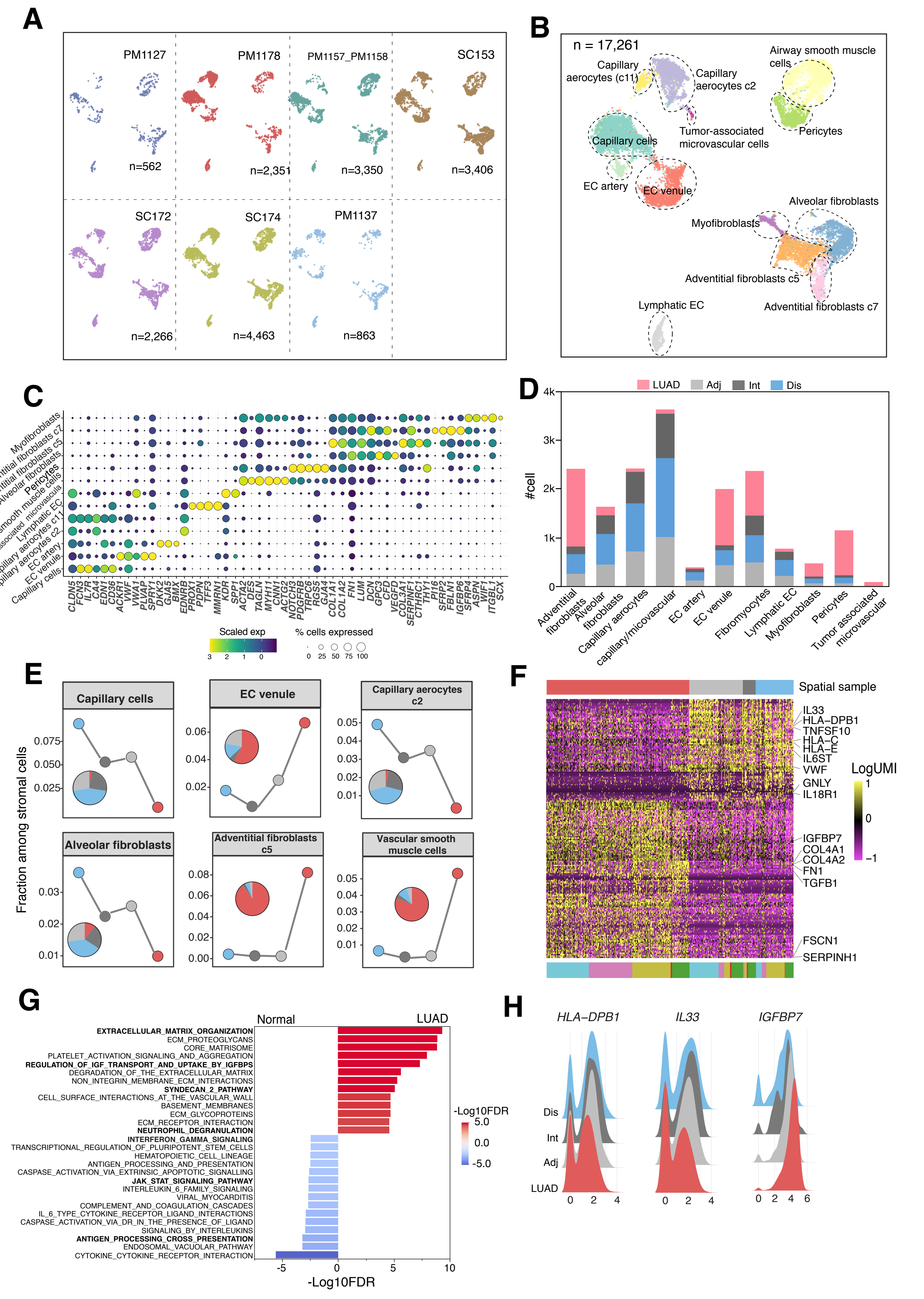

### Supplementary Fig. S21

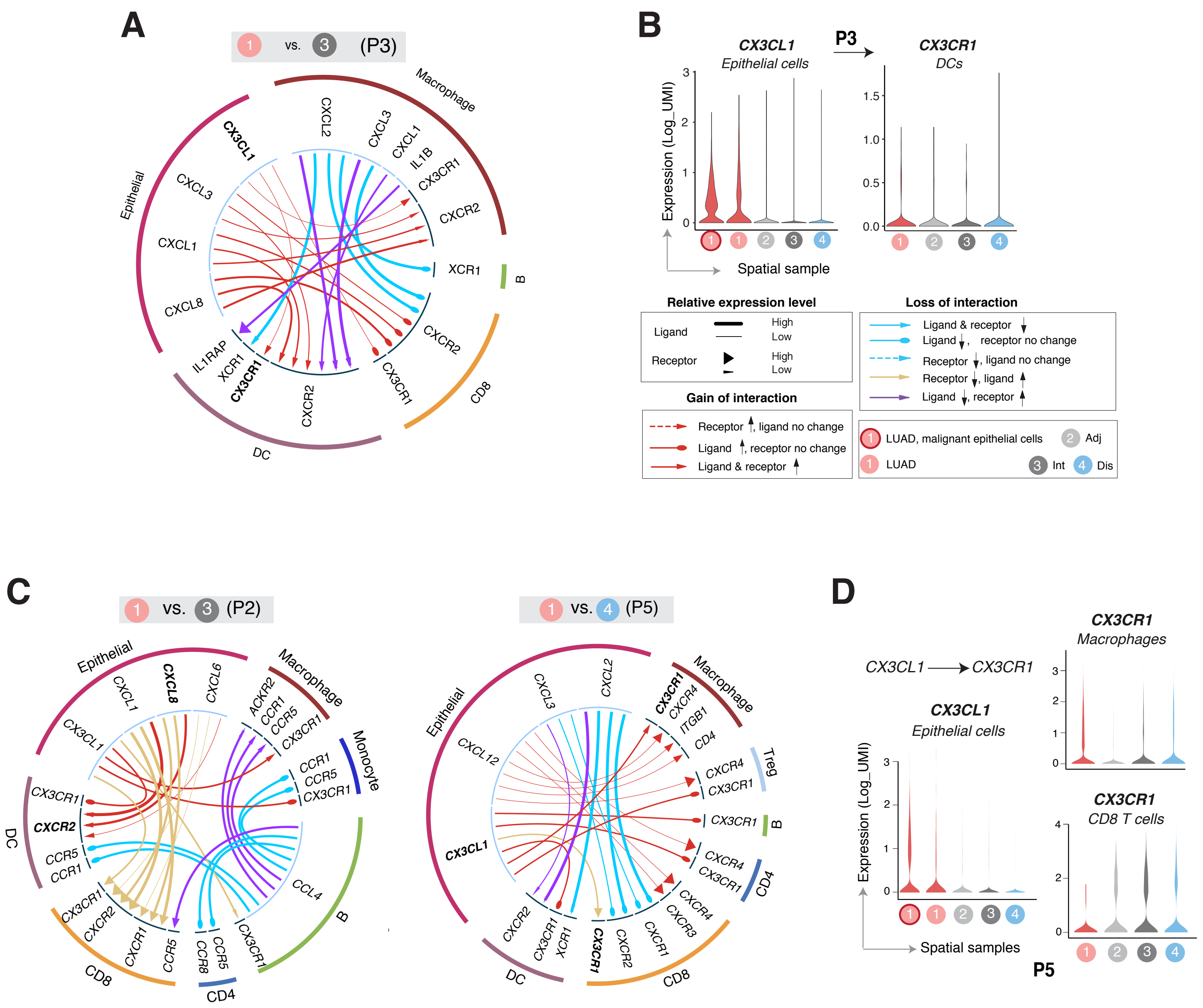

### Supplementary Fig. S22

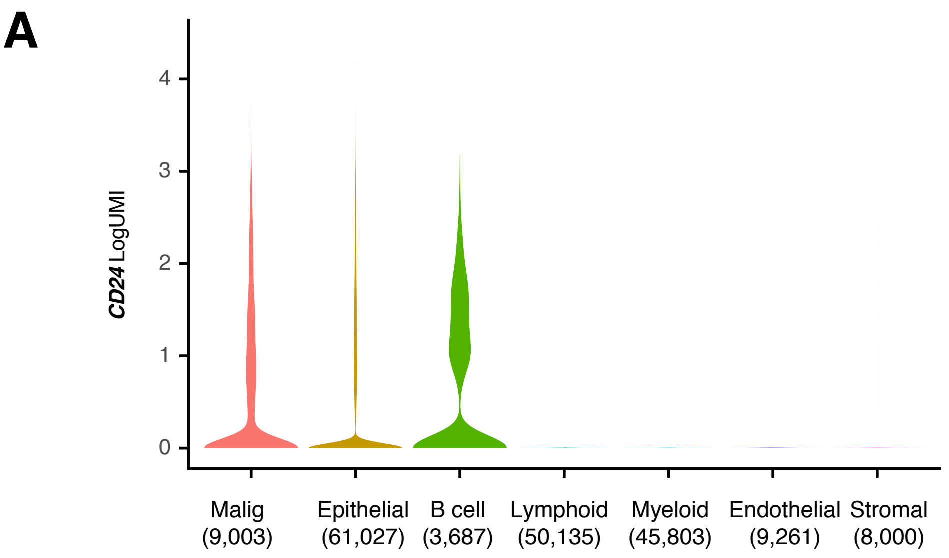


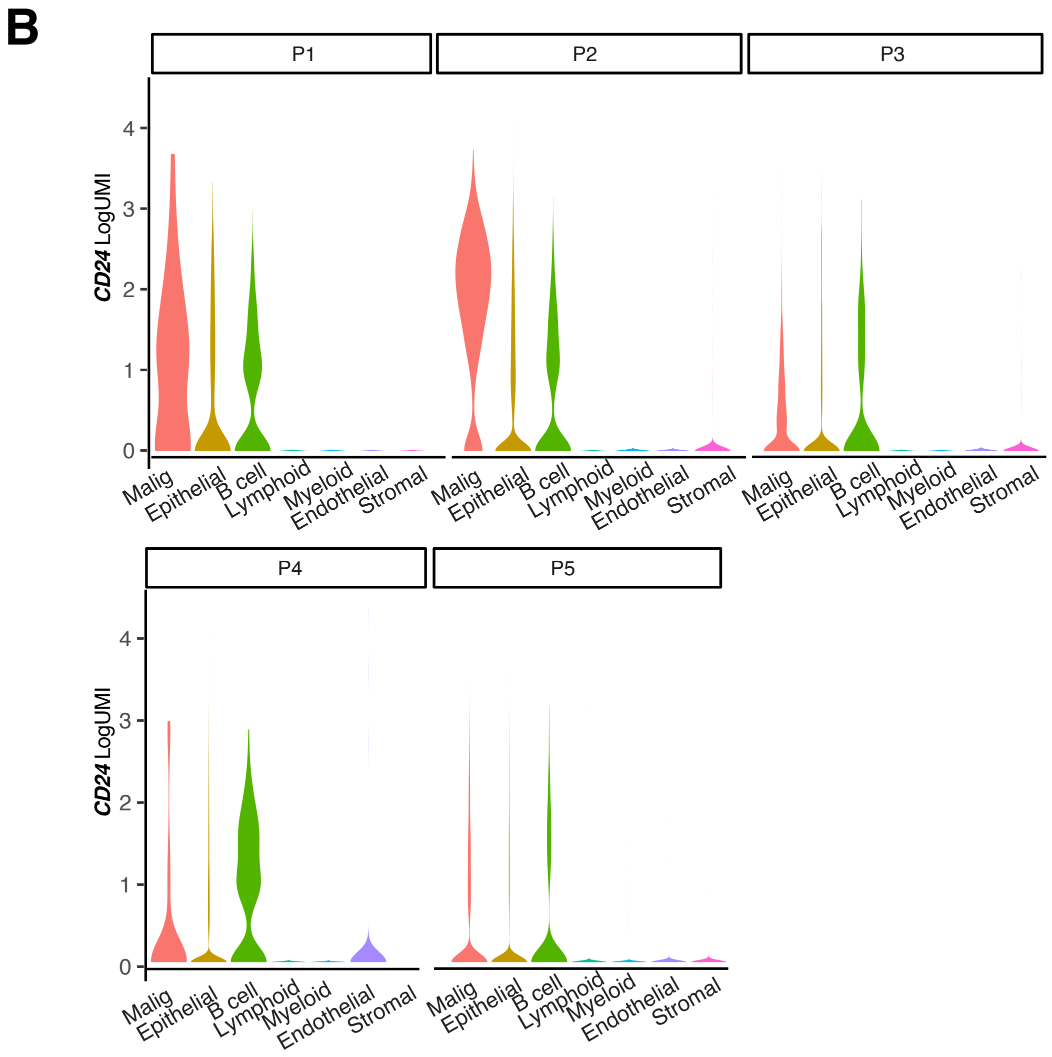

### Supplementary Fig. S23

## A

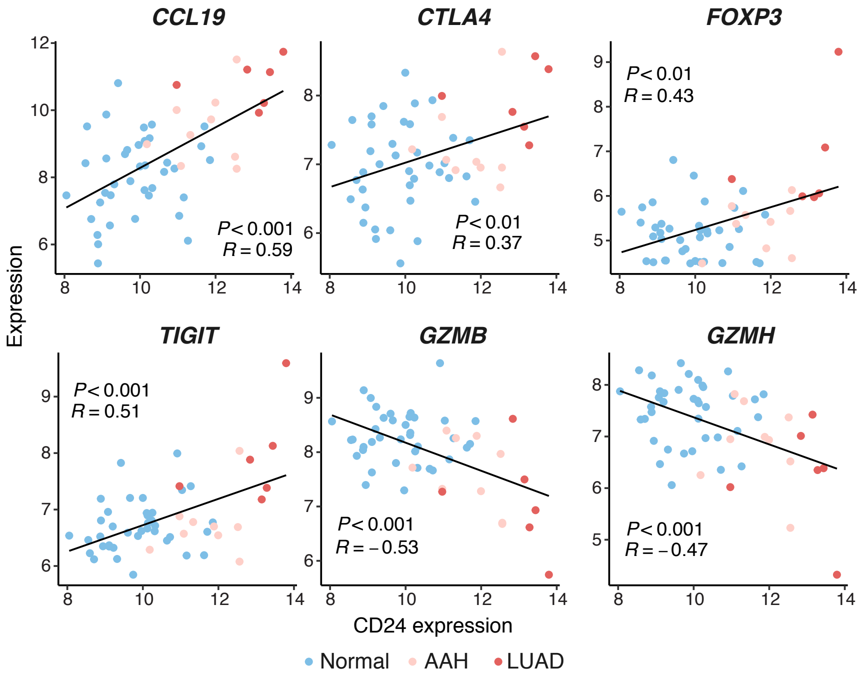


## B

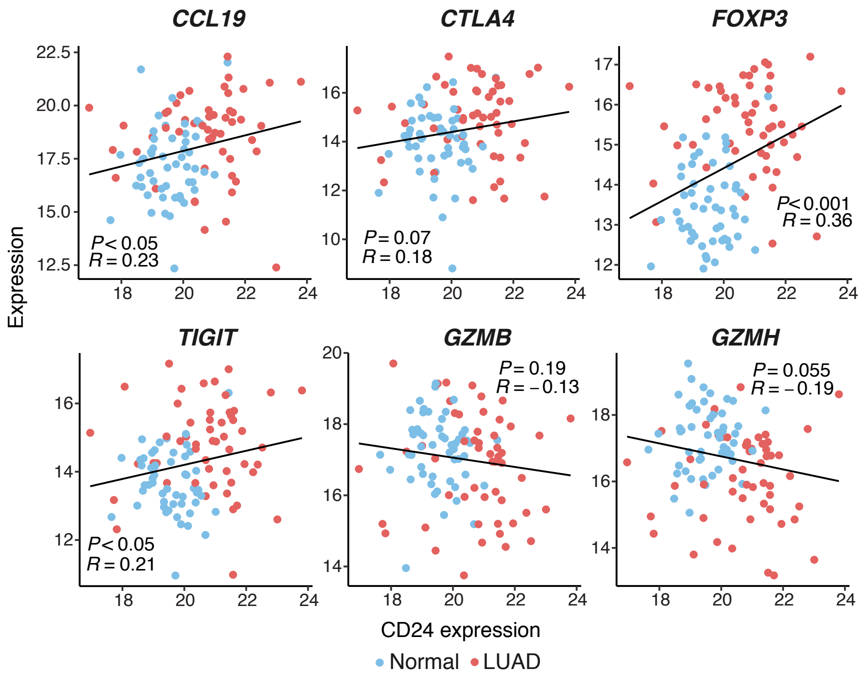

### Supplementary Fig. S24

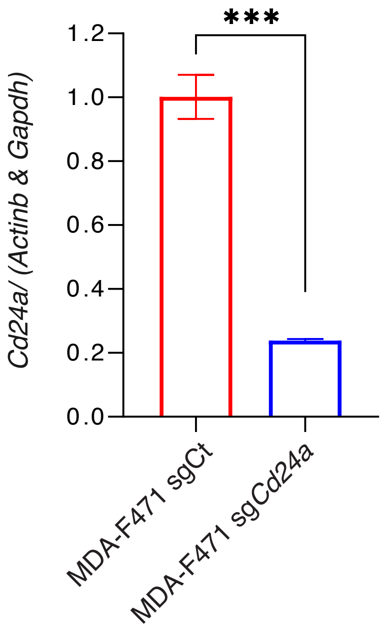
